# Antibody-Drug Conjugates Targeting the EGFR Ligand Epiregulin Elicit Robust Anti-Tumor Activity in Colorectal Cancer

**DOI:** 10.1101/2024.02.20.581056

**Authors:** Joan Jacob, Yasuaki Anami, Peyton High, Zhengdong Liang, Shraddha Subramanian, Sukhen C. Ghosh, Solmaz AghaAmiri, Cara Guernsey-Biddle, Ha Tran, Julie H. Rowe, Ali Azhdarinia, Kyoji Tsuchikama, Kendra S. Carmon

## Abstract

As colorectal cancer (CRC) remains a leading cause of cancer-related death, identifying therapeutic targets and approaches is essential to improve patient outcomes. The EGFR ligand epiregulin (EREG) is highly expressed in RAS wildtype and mutant CRC with minimal expression in normal tissues, making it an attractive target for antibody-drug conjugate (ADC) development. In this study, we produced and purified an EREG monoclonal antibody (mAb), H231, that had high specificity and affinity for human and mouse EREG. H231 also internalized to lysosomes, which is important for ADC payload release. ImmunoPET and *ex vivo* biodistribution studies showed significant tumor uptake of ^89^Zr-labeled H231 with minimal uptake in normal tissues. H231 was conjugated to either cleavable dipeptide or tripeptide chemical linkers attached to the DNA-alkylating payload duocarmycin DM, and cytotoxicity of EREG ADCs was assessed in a panel of CRC cell lines. EREG ADCs incorporating tripeptide linkers demonstrated the highest potency in EREG-expressing CRC cells irrespective of RAS mutations. Preclinical safety and efficacy studies showed EREG ADCs were well-tolerated, neutralized EGFR pathway activity, caused significant tumor growth inhibition or regression, and increased survival in CRC cell line and patient-derived xenograft models. These data suggest EREG is a promising target for the development of ADCs for treating CRC and other cancer types that express high levels of EREG. While the efficacy of clinically approved anti-EGFR mAbs are largely limited by RAS mutational status, EREG ADCs may show promise for both RAS mutant and wildtype patients, thus improving existing treatment options.

**Significance:** EREG-targeting antibody-drug conjugates demonstrate acceptable safety and robust therapeutic efficacy in RAS mutant and wildtype colorectal cancer, suggesting their potential as an alternative to EGFR-targeted therapy to benefit a broader patient population.

## Introduction

Colorectal cancer (CRC) ranks second in cancer-related deaths in the United States and incidence and mortality has considerably increased in individuals 50 years and younger (1). Chemotherapy combinations of irinotecan or oxaliplatin with fluoropyrimidines have long been considered first-line treatment options for advanced CRC, yet are limited by systemic toxicities (2). Approved monoclonal antibodies (mAbs) that target epidermal growth factor receptor (EGFR), cetuximab and panitumumab, have greatly improved overall survival for a fraction of patients with wildtype *KRAS* and *NRAS,* though ∼40-60% of patients present mutations (2). More recently, a phase 3 trial of *KRAS* G12C inhibitor, sotorasib, in combination with panitumumab showed superior progression-free survival rates versus standard of care for the 3-4% of metastatic CRC (mCRC) patients that harbor the specific mutation (3). Immune checkpoint inhibitors have shown durable response in tumors with microsatellite instability-high (MSI-H) mismatch repair deficiency, though the majority of mCRC is microsatellite stable (MSS) (2). While the development of targeted agents has improved prognosis for different CRC molecular subtypes, many patients still experience resistance and relapse, demonstrating the need for new targets and more effective therapeutic strategies.

Epiregulin (EREG), a member of the EGF family, is a surface-expressed, transmembrane ligand that binds and activates EGFR and human epidermal growth factor receptor 4 (HER4/ErbB4) and can stimulate signaling of HER2/ErbB2 and HER3/ErbB3 through receptor heterodimerization (4). EREG is highly expressed in many cancer types (5–9), including CRC (10), and has been shown to drive colitis-associated CRC (11) and be predictive of colorectal liver metastasis (12). EREG correlates with left primary tumor location (PTL), MSS, wildtype *BRAF* and *RAS* status, and CpG island methylator phenotype low (CIMP-low) CRC subtypes (10). Importantly, in RAS wildtype mCRC patients, EREG expression predicts benefit to EGFR-targeted therapies, including right PTL disease (13–17). Overall, EREG is a promising predictive biomarker for treatment and may serve as a novel therapeutic target.

Antibody-drug conjugates (ADCs) are an emerging class of anticancer drugs, consisting of highly specific mAbs chemically linked to potent cytotoxic payloads with low nanomolar to picomolar IC_50_ values (18,19). These armed mAbs directly deliver payloads to target antigen-enriched tumor cells while minimizing toxicity to normal tissues (20,21). Advances in linker-payload technology have improved ADC stability and safety profiles, leading to a surge of approved ADCs within the past few years and many more entering clinical trials (22). In this study, we generated and comprehensively characterized an EREG mAb for target-specific binding, internalization, species cross-reactivity, biodistribution, and tumor uptake using ^89^Zr-immunoPET. We then developed and evaluated a series of EREG-targeted ADCs incorporating diverse chemical linkers and the DNA minor groove alkylating payload duocarmycin DM (DuoDM). EREG ADCs were well-tolerated and demonstrated potent and selective growth inhibition in cancer cell lines and patient-derived tumor xenograft (PDX) models of CRC with or without mutant *RAS*. This preclinical data supports further development of EREG ADCs for treating EREG-high tumors irrespective of mutational status.

## Methods and Materials

### Plasmids and cloning

Sequences encoding human or mouse EREG (hEREG, Clone ID: 9020391 or mEREG, Clone ID: 5325124; Horizon Discovery) were subcloned and fused with the CD8 signal peptide sequence (MALPVTALLLPLALLLHAA) followed by a Myc-tag at the N-terminus in pIRESpuro3 (Clontech). Humanized EREG mAbs H231 and H206 (23) and E27 (24) were generated from patented sequences. The variable heavy and light chain (VH and VL) regions were synthesized (Epoch Life Science, Inc.), subcloned into pCEP4 containing human IgG1 heavy chain with N297A mutation for site-specific conjugation and kappa light chain constant regions. For H231, a Q57N mutation was incorporated in the VH region to prevent drug conjugation in the complementarity determining region. Similarly, cetuximab and CD20 mAb rituximab were generated utilizing publicly available sequences, the latter to serve as a non-targeting isotype control (cmAb). All expression constructs were sequence verified. For hEREG CRISPR-Cas9-based knockout (KO), single guide RNA sequence (sgRNA), 5’-GACAGAAGACAATCCACGTG-3’ was cloned into lentiCRISPRv2-hygro as described (25). LentiCRISPRv2-hygro was a gift from Brett Stringer (Addgene, #98291).

### Commercial antibodies and recombinant proteins

Primary antibodies used in this study include anti-β-actin (Cat# 4970, RRID:AB_2223172, 1:5000), anti-EREG (Cat# 12048, RRID:AB_2797808, 1:1000), anti-EGFR (Cat# 4267, RRID:AB_2864406, 1:1000), anti-phospho-EGFR Y1068 (Cat# 3777, RRID:AB_2096270, 1:1000), anti-AKT (Cat# 9272, RRID:AB_329827, 1:1000), anti-phospho-AKT S473 (Cat# 9271, RRID:AB_329825, 1:1000), anti-p44/42 MAPK/ ERK 1/2 (Cat# 9102, RRID:AB_330744, 1:1000), anti-phospho-p44/42 MAPK T202/Y204/ p-ERK 1/2 (Cat# 9101, RRID:AB_331646, 1:1000), anti-PARP (Cat# 9532, RRID:AB_659884, 1:1000), and anti-LAMP1 (Cat# 9091, RRID:AB_2687579, 1:400) from Cell Signaling Technology and anti-Myc-Cy3 (Sigma-Aldrich Cat# C6594 RRID:AB_258958, 1:1000). Recombinant hEREG (Cat# 1195EP025CF) was from R&D Systems.

### Antibody production

MAb production was performed by transiently expressing mAb constructs in Expi293F cells (ThermoFisher Cat# A14527, RRID:CVCL_D615) using polyethylenimine reagent (Polysciences, Cat# 247651). Medium was collected 7 days post-transfection and mAbs were purified using protein A resin (Genscript) affinity chromatography and eluted in PBS (pH 7.4). MAbs were analyzed for purity and homogeneity by SDS-PAGE and reverse-phase high-performance liquid chromatography (Vanquish UHPLC) and quantified by NanoDrop (ThermoFisher).

### Microbial transglutaminase (MTGase)-mediated antibody conjugation

For site-specific conjugation of Q295, H231 mAb (5.625 mL in PBS, 8.0 mg1mL , 45.0 mg) was incubated with diazide branched linker developed previously (26) (1801µL of 1001mM in H_2_O, 60 equiv.) and Activa TI (505 mg, final concentration 8%, Ajinomoto, Modernist Pantry) at RT for 20 h. The reaction was monitored using the Thermo LC-MS system consisting of a Vanquish UHPLC and Q Exactive™ Hybrid Quadrupole-Orbitrap™ Mass Spectrometer equipped with a MabPac RP column (2.1 × 50 mm, 4 µm, Thermo Scientific). Elution conditions were as follows: mobile phase A1= H_2_O (0.1% formic acid); mobile phase B1=1acetonitrile (0.1% formic acid); gradient over 3 min from A:B1=175:25 to 45:55; flow rate1=10.251mL1min . Conjugated mAb was purified by protein A column chromatography (MabSelect PrismA™, Cytiva) to afford a mAb-linker conjugate [42.3 mg, 94% yield determined by bicinchoninic acid (BCA) assay]. CmAb was conjugated similarly.

### Synthesis of DBCO-PEG3-EGC-PABQ-Duocarmycin DM-β-glucuronide

Fully protected DuoDM compound (26) (37.2 mg, 21.9 mmol) was dissolved in 40% TFA/DCM (600 mL and 800 mL). After being stirred at room temperature (RT) for 1 h, the solution was concentrated in vacuo and crude peptide was precipitated with cold diethyl ether (20 mL) followed by centrifugation at 2,000 ×g for 3 min (3 times). The crude compound was dissolved in MeOH (500 mL) and LiOH•H_2_O (23.7 mg, 564 mmol) in water (500 mL) was added to the mixture. After stirring at RT for 30 min, the mixture was cooled to 0°C and quenched with 6N-HCl/ACN. The solution was removed in vacuo and crude peptide was dissolved in DMF (500 mL). DBCO-NHS (9.1 mg, 22.6 µmol) and DIPEA (13 µL, 75.2 µmol) were added to the solution and the mixture was stirred at RT for 1 h. Crude products were purified by preparative RP-HPLC under basic conditions to afford analytically pure DBCO-PEG3-EGC-PABQ-DuoDM-β-glucuronide (EGC-qDuoDM gluc; 21.0 mg, 61% for the 3 steps). Purity was confirmed by LC-MS. White powder. HRMS (ESI) Calcd For C_79_H_91_N_11_O_21_Cl [M] : 1564.6074, Found: 1564.6072.

### Strain-promoted azide-alkyne cycloaddition to attach payload

The EGC-qDuoDM gluc was added (76.4 µL of 10 mM stock solution in DMSO, 1.5 equivalent per azide) to a solution of H231-or cmAb-linker conjugate in PBS (4.71mL, 4.0651mg mL–1), and the mixture was incubated at RT for 30 min. The reaction was monitored using the Thermo LC-MS system with elution conditions stated previously. The crude products were purified by CHT-XT (Bio-Rad) after exchanging buffer to 5 mM phosphate buffer pH 6.5. Desired ADC was eluted by salt gradient and exchanged buffer to PBS. Yield was determined 79% (15.14 mg). Drug-antibody ratio (DAR) values for H231 EGC-qDuoDM gluc and cmAb EGC-qDuoDM gluc (cADC) were determined by Electrospray ionization-mass spectrometry (ESI-MS) analysis. Other ADCs incorporating BCN-PEG3–EGCit–PABC–DuoDM (EGC-cDuoDM) or DBCO-PEG4-VC-PAB-(PEG2)-DuoDM (VC-cDuoDM; Levena) were prepared in a similar manner or according to our previous reports (26).

### Cell culture and transfection

DLD-1 (Cat# CCL-221, RRID:CVCL_0248), LoVo (Cat# CCL-229, RRID:CVCL_0399), RKO (Cat# CRL-2577, RRID:CVCL_0504), SW48 (Cat# CCL-231, RRID:CVCL_1724), LS180 (Cat# CL-187, RRID:CVCL_0397), HCT116 (Cat# CCL-247, RRID:CVCL_0291), COLO320 (CCL-220, RRID:CVCL_0219), HCT15 (Cat# CCL-225, RRID:CVCL_0292), HT-29 (Cat# HTB-38, RRID:CVCL_0320), SW403 (Cat# CCL-230, RRID:CVCL_0545), SW620 (Cat# CCL-227, RRID:CVCL_0547), and HEK293T/293T (Cat# CRL-3216, RRID:CVCL_0063) cells were purchased from ATCC. LIM1215 (Cat# 10092301, RRID:CVCL_2574) were purchased from Millipore Sigma. Most cell lines were authenticated utilizing short tandem repeat profiling and tested for mycoplasma prior to 2019 and were passaged less than 15 times, with SW403 and LIM1215 passaged less than 5 times and not further authenticated due to ATCC purchase after 2020. 293Ts were cultured in DMEM and CRC cells in RPMI medium supplemented with 10% fetal bovine serum and penicillin/streptomycin. LIM1215 cells were additionally supplemented with 1 µg/ml hydrocortisone, 10 µM 1-Thioglycerol, 0.6 µg/ml insulin and 25 mM HEPES. Cell lines were cultured at 37°C with 95% humidity and 5% CO_2_. To generate the DLD-1 EREG KO cell line, lentivirus particles were produced by co-transfecting 293Ts with lentiCRISPRv2-hygro incorporating EREG sgRNA and packaging plasmids, psPAX2 and pMD2.G, using Fugene HD (Promega Cat# E2311). Clonal selection was performed using 50 µg/ml hygromycin.

### Western blot

Protein extraction was performed using RIPA buffer supplemented with protease/phosphatase inhibitors. PDX tissues were homogenized and freeze thawed. Lysates were centrifuged 15 min, 4 °C at 13,000 x g. Protein concentrations were measured and lysates were diluted in Laemmli buffer and incubated at 37°C for 1 h prior to running SDS-PAGE. HRP-labeled secondary antibodies were utilized with the standard ECL protocol. For EREG mAb neutralization experiments, cells were seeded in a 6-well plate and cultured in 0.5% FBS in RPMI 3 h then treated with PBS, 300 ng/ml recombinant hEREG, 15 µg/ml cmAb or H231 mAb, or combination for 5 min. For PARP analysis, cells were treated with H231 EGC-qDuoDM gluc or cADC as indicated.

### Cell-based binding assays

293T cells stably overexpressing hEREG, mEREG, or vector or CRC cells were seeded onto poly-D-lysine-coated black 96-well plates and incubated overnight. Serial dilutions of H231, cmAb, H231 ADCs, or cADC were added for 2 h at 4°C. Plates were washed in PBS, fixed in 4% formalin, and incubated with anti-human-Alexa-555 (ThermoFisher Cat# A-21433, RRID:AB_2535854) for 1 h at RT. Fluorescence intensity was quantified using Tecan Infinite M1000 plate reader (RRID:SCR_025732). Each condition was tested in triplicate in 2-3 independent experiments.

### Flow cytometry

Surface-expressed EREG molecules/cell were quantified by labeling H231 with phycoerythrin (PE) with the Lightning-Link PE conjugation kit (Abcam Cat# ab102918) and BD QuantiBRITE PE (BD Biosciences Cat# 340495). Briefly, 1×10 cells/sample were fixed with dropwise addition of 10% formalin for 20 min, followed by blocking with 2% BSA/PBS for 5 min and incubation with 6 µg/ml H231 or cmAb in 2% BSA/PBS at 4°C for 60 min. QuantiBRITE PE beads were run on BD FACS ARIA II SORP flow cytometer to generate the standard curve for fluorescent intensity vs. number of PE molecules/bead. Singlets were gated on the FSC-H vs. SSC plot, and the singlet bead population was analyzed using a histogram plot of FL-2 axis in linear values. Log10 for lot-specific PE/bead values vs. Log10 of geometric means for four bead peaks were plotted. Equation y=mx+c where x equals Log10 PE molecules/bead and y equals Log10 fluorescence was used to calculate slope, intercept and correlation coefficient. To determine mAbs bound per cell for different cell lines, samples were run using same settings and PE molecules/cell were calculated. Unstained cells and cmAb served as negative controls.

### Immunocytochemistry

CRC cells were seeded into poly-D-lysine coated 8-well culture slides and incubated overnight. Cells were treated with H231 at 37°C for 90 mins to allow binding and internalization, then washed, fixed in 4% formalin, permeabilized in 0.1% saponin, incubated with anti-LAMP1 at RT for 45 min, followed by anti-rabbit-Alexa-488 (ThermoFisher A-11008, RRID:AB_143165) and anti-human-Alexa-555 (ThermoFisher Cat# A-21433, RRID:AB_2535854) at RT for 1 h. Nuclei were counterstained with TO-PRO-3 (ThermoFisher Cat# T3605). Images were acquired using confocal microscopy (Leica TCS SP5, RRID:SCR_020233) and LAS AF Lite software. Co-localization was quantified using ImageJ (RRID:SCR_003070).

### ^89^Zr-immunoPET and biodistribution

Immuno-positron emission tomography (immunoPET) studies were carried out in accordance with IACUC at UTHealth Houston (AWC-20-0144 and AWC-23-0106). 2.5 x 10 DLD-1 cells in 50 % Matrigel were subcutaneously implanted into the lower right flank of female 6-8-week-old nu/nu mice.

Desferrioxamine (DFO) immunoconjugates were prepared via lysine conjugation as previously described (27). DFO-H231 or DFO-cmAb (200µg) were radiolabeled with zirconium-89 (^89^Zr; Washington University, St. Louis, MO). Once tumors reached ∼4-6 mm diameter, Zr-H231 or ^89^Zr-cmAb were injected with 139-215 µCi per mouse. Imaging was performed 5 days post-injection (dpi) using an Albira small animal PET/CT scanner (Bruker) and tumor-to-muscle ratios where determined. Mice were euthanized and organs were resected, weighted, and analyzed using a Cobra II auto-γ counter (Packard) to determine tracer biodistribution. Standard aliquots of known activity were used to calculate the percentage of the injected dose per gram of tissue (%ID/g).

### In vitro viability

CRC cells were plated at ∼1,000 cells/well in 96 half-well plates. Serial dilutions of unconjugated H231, cmAb, H231 ADCs or cADC were added and incubated at 37 °C for 4 or 5 days. Cell viability was measured using CellTiter-Glo 2.0 (Promega Cat# G9242) according to the manufacturer’s protocol. Luminescence was measured using Tecan Infinite M1000 plate reader (RRID:SCR_025732).

### Tolerability Study

For single-dose studies, female 6-8-week-old C57BL/6J mice (n=14/group, The Jackson Laboratory, RRID:IMSR_JAX:000664) received a single dose H231 VC-DuoDM (5 or 10 mg/kg), H231 EGC-cDuoDM ADC (5 or 10 mg/kg), or PBS vehicle via tail vein. For multiple-dose studies, female 6-8-week-old C57BL/6 mice (n=13-4/group) were administered 5 mg/kg H231, cADC, or H231 EGC-qDuoDM gluc, or vehicle weekly for 3 weeks. Body weight was monitored every 2-3 days for 2 weeks after treatment termination. Humane endpoints were defined as >20% weight loss or severe signs of distress. However, no mice met these criteria throughout the study. Fourteen dpi, mice were terminally anesthetized and whole blood was drawn by heart puncture for blood chemistry analysis in the Department Veterinary Medicine & Surgery at MD Anderson Cancer Center. Liver and kidney tissues were collected and then paraffin embedded, sectioned and H&E stained in the histopathology core at UTHealth Houston Institute of Molecular Medicine.

### In vivo efficacy studies

Animal studies were carried out in accordance with IACUC (AWC-20-0144, AWC-21-0142, and AWC-23-0106). For efficacy studies, nu/nu (Charles River Laboratories, RRID:IMSR_CRL:088 or The Jackson Laboratory, RRID:IMSR_JAX:002019) and NSG (The Jackson Laboratory, RRID:IMSR_JAX:005557) mice were used for CRC cell xenografts and PDX models, respectively. PDX CRC-001 was from Jackson Laboratory (#TM00849) and XST-GI-010 was established in our laboratory at UTHealth with patient consent and appropriate approvals (HSC-MS-20-0327, HSC-MS-21-0074). Female 6-8-week-old mice were implanted subcutaneously with 3-4 x 10 LoVo or DLD-1 CRC cells or 2-3 mm PDX fragments into the right flank. When tumors reached ∼100–200 mm , mice were randomized based on equivalent average tumor volume for each treatment group. Animals without palpable tumors were excluded. No blinding to group allocation was performed and sample sizes were determined based on prior experience with the models to reach statistical significance. Mice were intravenously or interperitoneally dosed weekly with PBS vehicle, H231, cADC, H231 ADCs, or cetuximab as indicated. Mice were routinely monitored for morbidity and mortality. Tumor volumes were measured bi-weekly and estimated by the formula: tumor volume = (length × width)/2. Mice were euthanized when tumor diameter reached 15 mm. Percentage of tumor growth inhibition (% TGI) was calculated using the formula [1 – (change of tumor volume in treatment group/change of tumor volume in vehicle group)] × 100 (%).

### Statistical Analysis

All data were analyzed using GraphPad Prism software (RRID:SCR_002798). Data are presented as mean1±1SEM or1±1SD, as indicated. In vitro binding and viability data were analyzed using the logistic nonlinear regression model. Each condition was tested in triplicate in 3 independent experiments. The Cancer Genome Atlas (TCGA) data from matched normal and tumor samples or *KRAS* status were compared and analyzed using paired Student’s t test. EREG in tumor vs. normal tissues and distribution in MSI/MSS was compared using one-way analysis of variance (ANOVA) and Tukey’s multiple comparison test. Kaplan-Meier analysis was performed to determine significance of EREG on disease-free and overall survival. Statistical significance of in vitro experiments was analyzed with ANOVA and Tukey’s multiple comparison test. Differences between groups for biodistribution studies were analyzed by Student’s t test. Statistical analysis of tumor volume was assessed by one-way ANOVA and Dunnett’s multiple comparison test or Student’s t test for two groups. P value less than < 0.05 was considered significant.

### Data and Materials Availability

The data generated in this study are available within the article and supplementary files. Materials and reagents will be made available through collaboration in agreement with UTHealth Houston and the corresponding author.

## Results

### EREG is highly expressed in MSS colorectal tumors with minimal expression in normal tissues

We analyzed *EREG* expression in TCGA RNA-seq data to identify CRC patients that may benefit from EREG-targeted ADCs (28). Though *EREG* has minimal expression in normal tissues, mRNA can be detected in skin, esophagus, vagina, lung, and colon as confirmed the Genotype-Tissue Expression (GTEx) project (Fig. 1A and Supplementary Fig. S1). Importantly, *EREG* is significantly higher in tumors from the colorectal adenocarcinoma (COADREAD) cohort compared to normal colon/rectum, lung, esophagus (average fold differences ≈ 3.7, 7.1, and 9.6, respectively; Fig. 1A) and other tissues. Matched tumor and adjacent normal samples from 32 patients showed *EREG* is upregulated >2-fold in ∼60% of patients (average fold difference in EREG-high patients ≈ 16.8; Fig. 1B). *EREG* expression is significantly upregulated in MSI-L and MSS tumors compared with MSI-H tumors (Fig. 1C), consistent with previous reports from other CRC cohorts (10,17). Furthermore, *EREG* levels are highly expressed in subsets of both *KRAS* wildtype and mutant (G12/G13) patients, however there is significant association with wildtype status (Fig. 1D). Partitioning of data from the COADREAD cohort showed no significant correlation between *EREG* expression and survival (Figs. 1E-F), though EREG-high patients trended toward poorer survival at later time-points (disease-free, median value = high, 63.37 vs. low, 78.65 months; overall survival, median value = 65.80 vs. 92.67 months). Notably, EREG-high status confers an overall survival advantage at earlier time-points (Fig. 1F), which has been shown to be significant for KRAS wildtype patients who received anti-EGFR therapy (17,29). Analysis of a recent study of 18 matched primary tumors, synchronous liver metastases, and normal colonic epithelium (17 MSS and 1 MSI-H) showed *EREG* was significantly elevated in both primary tumors and their metastases (Fig. 1G) (30). Western blot analysis of EREG in MSS, *RAS* mutant CRC PDX models showed high EREG expression in 4/5 samples (Fig. 1H). Furthermore, RNA-seq data from the Cancer Cell Line Encyclopedia (CCLE) revealed *EREG* is amply expressed in a majority of CRC cell lines irrespective of mutant *RAS* and was consistent with protein levels (Fig. 1I-J) (31). These results show EREG correlates with MSI-L/MSS in patient tumors, which accounts for the majority of mCRC patients, and is highly expressed in a fraction of both *RAS* wild-type and mutant subtypes, suggesting EREG ADCs may be a promising treatment option for a sizeable patient population.

**Figure 1.**
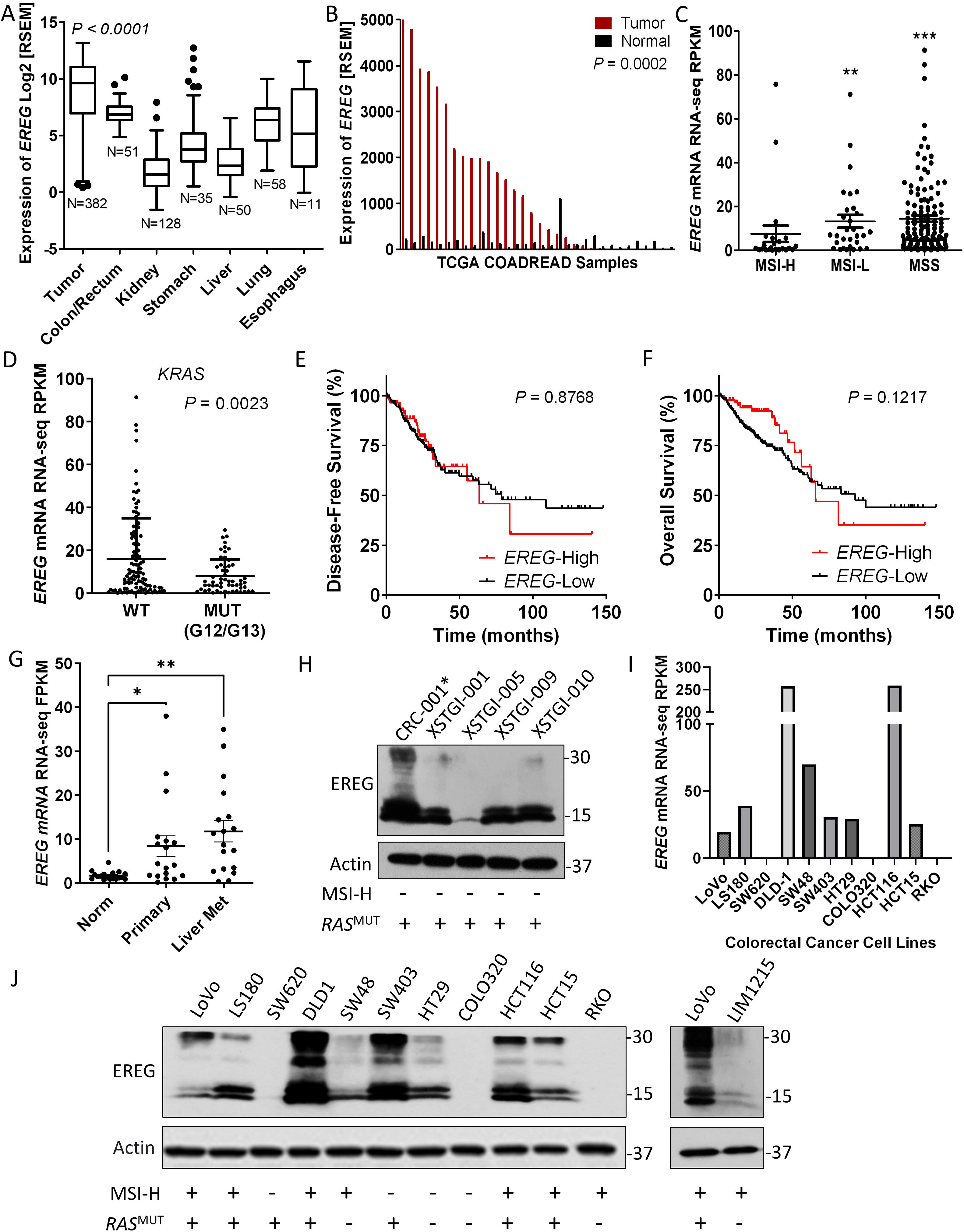
EREG is highly expressed in colorectal tumors and cancer cell lines. **A** , EREG RNA-Seq expression data from patient tumors of the colorectal adenocarcinoma (COADREAD) TCGA cohort. Values are Log2 RNA-Seq by expectation–maximization (Log2 RSEM). **B**, EREG RNA-Seq RSEM values for matched tumor and adjacent normal from the COADREAD cohort (N=32). Distribution of **C**, MSI-H (N=23), MSI-L (N=30), and MSS (N=129) subtypes and **D**, KRAS WT (N=109) and MUT (N=56) status based on EREG expression. Values are read per kilobase of transcript per million (RPKM). Kaplan-Meier analyses of **E**, disease-free survival (EREG-High, N=86; EREG-Low, N=245) and **F**, overall survival (EREG-High, N=92; EREG-Low, N=279) in patients from the COADREAD cohort. P values were obtained by the log-rank test. **G**, EREG RNA-seq values from cohort of 18 matched normal colonic epithelium, primary adenocarcinoma, and synchronous liver metastases (NCBI GSE50760). **H**, Western blot endogenous EREG protein expression in patient-derived xenograft (PDX) models. *, indicates mutant NRAS. **I**, EREG RNA-seq values for select CRC cell lines used in this study from Cancer Cell Line Encyclopedia (CCLE). **J**, Western blot of endogenous EREG protein expression across CRC cell lines. Quantitative data presented as mean ± SEM and statistical analysis performed using one-way analysis of variance (ANOVA) and Tukey’s multiple comparison test or paired Student’s t test for two groups. ** P< 0.01, *** P< 0.001.

### H231 binding and internalization in CRC cell lines

To develop a novel EREG-targeted ADC, we first set out to identify an optimal EREG mAb with high affinity and specificity. We cloned and purified three individual EREG mAbs that bind both surface-expressed, uncleaved pro-EREG and mature, cleaved EREG: H206, H231, and E27 (23,24). To determine mAb target specificity, we utilized 293T cells that do not express endogenous EREG and established lines overexpressing myc-tagged hEREG, mEREG, or vector. Expression was confirmed using anti-myc-tag antibody (Supplementary Fig. S2A). Adherent cell-based binding assays show all EREG mAbs bind hEREG-293T cells (Kd values = H231, 0.01 µg/ml or 0.1 nmol/L; H206, 0.06 µg/ml or 0.4 nmol/L; E27, 0.08 µg/ml or 5.6 nmol/L; Fig. 2A and Supplementary Fig. S2B). H206 (Kd = 0.10 µg/ml or 0.7 nmol/L) and H231 (Kd = 0.18 µg/ml or 1.2 nmol/L), but not E27, bind mEREG-293T cells (Fig. 2A and Supplementary Fig. S2C). Non-targeting cmAb shows no binding and H231 failed to bind vector cells, verifying specificity (Fig. 2A and Supplementary Fig. S2C). H231 was selected as lead mAb for further characterization and ADC development due to its high affinity and maximum binding, species cross-reactivity, and high production yield. We showed H231 has high affinity binding to endogenous EREG on DLD-1 (Kd= 0.36 µg/ml or 2.4 nmol/L; Fig. 2B) and LoVo (Kd= 0.15 µg/ml or 1.0 nmol/L; Fig. 2C) cells. No binding was detected for cmAb. We next quantified surface pro-EREG ligands/cell. CRC cells were incubated with H231 or cmAb conjugated with PE or left unstained and flow cytometric analysis was performed. DLD-1 EREG-KO cell line was established using CRISPR-Cas9 to further verify H231 specificity for endogenous EREG. KO was confirmed by western blot (Fig. 2D). Pro-EREG ranged between 218-23,937 ligands/cell in EREG-high cells (HCT116 and DLD-1) and was much lower in CRC cells with moderate (SW48) to low (SW620) EREG expression and DLD-1 EREG-KO cells (Fig. 2E and Supplementary Table S1). Mean pro-EREG/cell for HCT116 and DLD-1 cells were 4,988 and 9,103, respectively, consistent with the acceptable number of targets/cell for clinically approved ADCs (Fig. 2E and Supplementary Table S1) (32,33).

**Figure 2.**
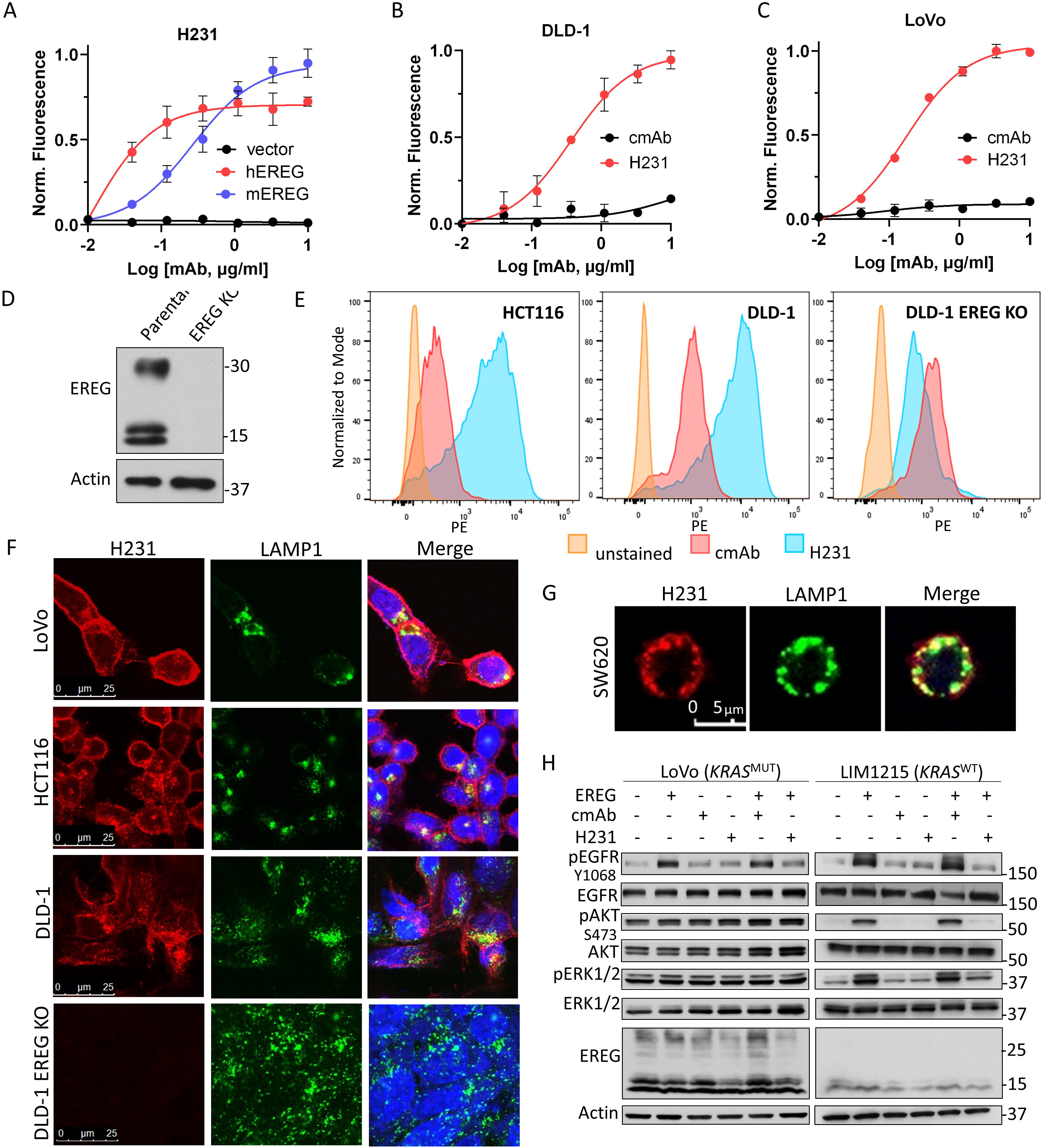
Characterization of H231 mAb binding, neutralization, and internalization. **A** , Cell-based binding assays show H3211mAb binds recombinant hEREG-and mEREG-overexpressing 293T cells and not vector cells. H231 and not cmAb binds endogenous EREG in **B**, DLD-1 and **C**, LoVo cells. Data presented as mean ± SD. **D** , Western blot of DLD-1 cells with high EREG expression stably transduced with CRISPR-Cas9 vector containing sgRNA to generate EREG knockout (KO) line. **E**, H231 and cmAb conjugated to phycoerythrin (PE) were utilized for the flow cytometric analysis to quantify surface pro-EREG ligands/cell using QuantBRITE PE kit. **F**, Confocal images shows H231 binds to EREG and co-localizes with lysosome marker, LAMP1, after 90 mins at 371°C in EREG-expressing LoVo, HCT116, and DLD-1 cells and not DLD-1 EREG KO cells. Yellow indicates colocalization. **G**, Overexpressed (O.E.) EREG co-localizes with lysosomes in EGFR-negative SW620 CRC cells after H231 treatment. **H**, Western blot of LoVo and LIM1215 cells treated in the presence of 15 µg/ml cmAb or H231 for 5 min with or without 300 ng/ml EREG.

Along with cell surface availability, an ideal target for ADC development should rapidly internalize and traffic to the lysosome for payload release. Immunocytochemistry (ICC) data shows H231 binds EREG-expressing LoVo, HCT116, and DLD-1 cells, internalizes, and co-localizes with the lysosome marker, LAMP1, after 90 mins at 37 °C (Fig. 2F). Notably, quantification of H231 time-course treatment verified peak co-localization occurs at 90 min (Supplementary Fig. S2D). As anticipated, H231 does not bind DLD-1 EREG KO cells and cmAb does not bind any CRC cell lines (Fig. 2F and Supplementary Fig. S2E). Next, to determine if EREG internalization is dependent on the presence of its receptors, we overexpressed hEREG in SW620 cells that do not express endogenous EREG, EGFR, or HER4/ERBB4 (Fig. 1I and Supplementary Fig. S2F). As shown in Fig. 2G, the H231-EREG complex effectively co-internalized to lysosomes, suggesting H231-mediated EREG internalization can occur independent of EGFR and HER4.

### H231 Neutralization Activity

Mature, cleaved EREG binds to EGFR to potentiate signaling through PI3K/AKT or MAPK pathways (4). Thus, we tested the ability for H231 to neutralize EREG signaling. *KRAS* mutant LoVo and DLD-1 cells, the latter which also harbor a *PIK3CA* mutation, and *KRAS* wildtype LIM1215 cells were treated with recombinant hEREG in the presence and absence of H231 or cmAb. Treatment with hEREG for 5 min increased phosphorylation of EGFR in LoVo and LIM1215 cells, with minimal effects in DLD-1 cells (Fig. 2H and Supplementary Fig. S2G). Phosphorylation of ERK1/2 and AKT was detected in LIM1215 cells and, interestingly, hEREG induced p-ERK1/2 in DLD-1 cells despite its mutational status (Fig. 2H and Supplementary Fig. S2G). Importantly, addition of H231, and not cmAb, inhibited downstream signaling mediated by hEREG, demonstrating H231-mediated neutralization (Fig. 2H).

### ImmunoPET and biodistribution of ^89^Zr-labeled H231

ImmunoPET imaging using ^89^Zr-labeled mAbs provides a means to measure whole-body biodistribution and verify optimal tumor delivery of therapeutic mAbs to predict safety and efficacy (20,27). First, H231-DFO and cmAb-DFO immunoconjugates were generated and HPLC showed minimal variance in retention time and peak profiles, with no effect on H231 affinity (Fig. 3A and Supplementary Figs. S3A-B). The Zr-labeled immunoconjugates were produced with radiochemical yields and purities of >70% and >99%, respectively (Supplementary Fig. S3C). PET imaging of EREG-high DLD-1 cell xenografts demonstrated higher tumor uptake with ^89^Zr-H231 compared to ^89^Zr-cmAb, which showed minimal on-target accumulation 5 days after injection (Fig. 3B). As shown in Fig. 3C, tumor-to-muscle ratios obtained from the PET images were nearly 5-fold higher for ^89^Zr-H231 compared to the control agent (12.6 ± 6.3 vs 2.7 ± 0.4). Ex vivo biodistribution of tumor and normal tissues revealed 3.8-fold higher tumor uptake for 89Zr-H231 with no notable EREG-specific uptake in other organs. Both tracers showed the expected clearance through the liver, while higher renal clearance was seen with Zr-H231 (Fig. 3D).

**Figure 3.**
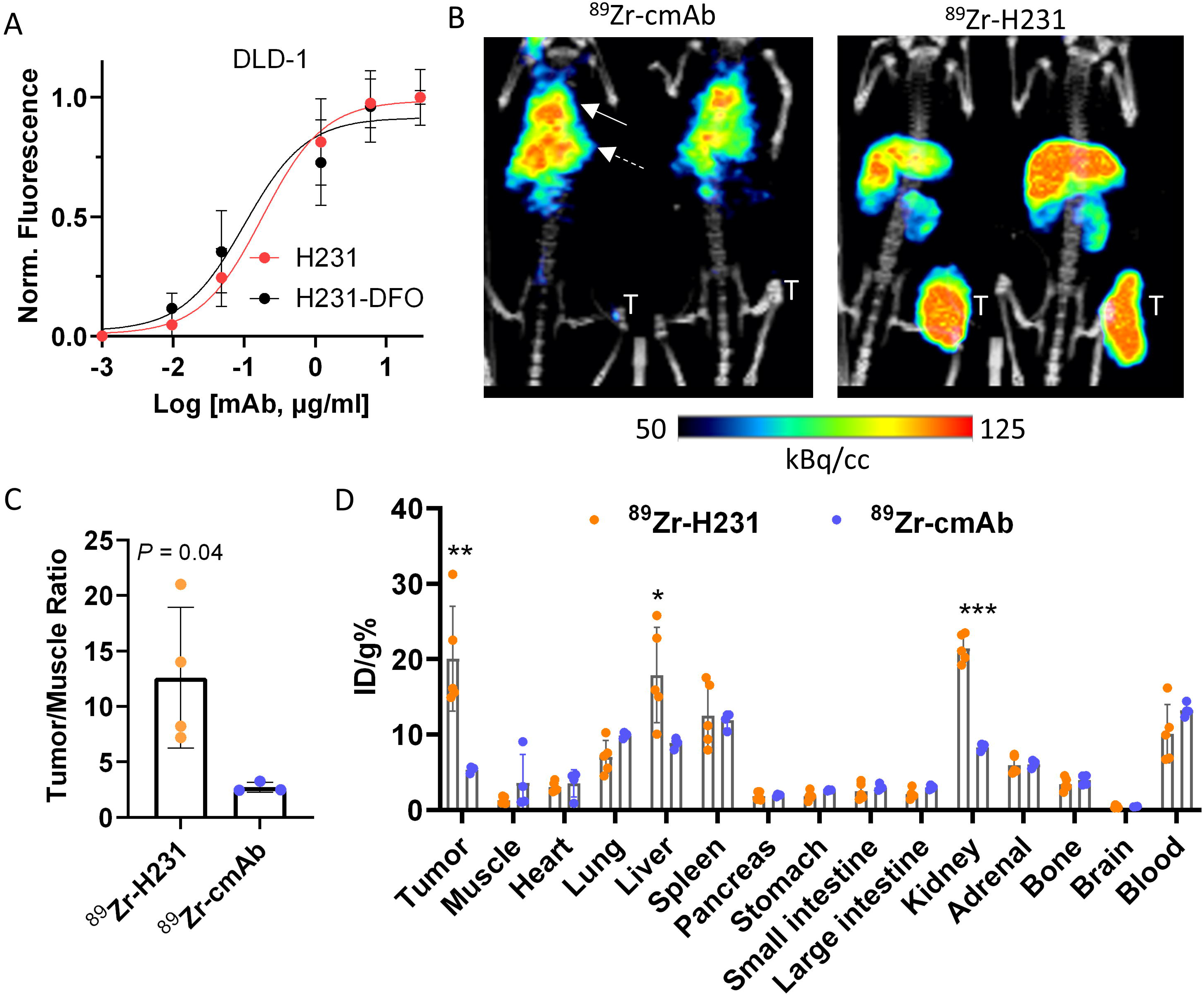
ImmunoPET imaging and biodistribution of H231 in EREG-high colorectal tumors. **A**, Cell-based binding assay shows equivalent binding for unconjugated H231 mAb and lysine-conjugated H231-DFO on DLD-1 cells. **B,** Representative PET/CT images acquired 5 days post-injection of ^89^Zr-radiolabeled EREG-targeting H231 mAb or non-targeting cmAb in DLD-1 tumor xenografts. Solid and dashed arrows indicate the lung and liver, respectively. **C,** Quantification of PET images tumor/muscle ratio (%ID/cc) of ^89^Zr-H231 (N=4) and ^89^Zr-cmAb (N=3). **D,** Biodistribution data (ID/g%) measured ex vivo in tumor and normal tissues ( Zr-H231, N=5; Zr-cmAb (N=4). Statistical analysis using Student’s t-test. Quantitative data presented as mean ± SD. \**P*< 0.05, \*\**P*< 0.01, *** *P*< 0.001.

### Generation and validation of EREG ADCs using diverse linker-duocarmycin payloads

For development of an optimal EREG ADC, we generated three candidates with diverse linkers incorporating highly cytotoxic duocarmycins, which we previously showed to be an effective payload in CRC (34). H231-based ADCs were produced via a site-specific chemo-enzymatic approach using MTGase as previously described (Fig. 4A) (35). Briefly, a branched bis-azido linker was conjugated to H231 via the Q295 sites of the Fc region, followed by strain-promoted alkyne-azide cycloaddition to attach a series of linker-duocarmycin DM (DuoDM) payloads incorporating clinically approved dipeptide (valine-citrulline; VC) or novel tripeptide (glutamic acid-glycine-citrulline; EGC) cleavable linkers shown to enhance ADC hydrophilicity and stability (Fig. 4A and Supplementary Fig. S4A-C) (26). Successful production and purification of EGC-cDuoDM was previously shown (26) and novel EGC-qDuoDM gluc linker-payload showed product purity >95% (Supplementary Fig. S5). Importantly, β-glucuronidation of qDuoDM-gluc allows it to function more as a prodrug that is more hydrophilic compared to cDuoDM (36). ESI-MS analysis confirmed homogeneity of each ADC with a uniform DAR= 4 (Fig. 4B) and Coomassie SDS-PAGE and size-exclusion chromatography (SEC) revealed high purity with no significant aggregation or dissociation (Supplementary Figs. S6 and S7A). Non-targeting cADC was generated with EGC-qDuoDM gluc and validated similarly (Supplementary Fig. S7B). Of note, we also attempted to produce similar H231 ADCs using mouse in place human antibody framework. However, further engineering is required due to additional Q sites in the mouse IgG1 Fc region accessible to MTGase conjugation. Linker-payload conjugation did not affect H231 binding to LoVo and DLD-1 cells and cADC showed no binding (Supplementary Fig. S7C-D).

**Figure 4.**
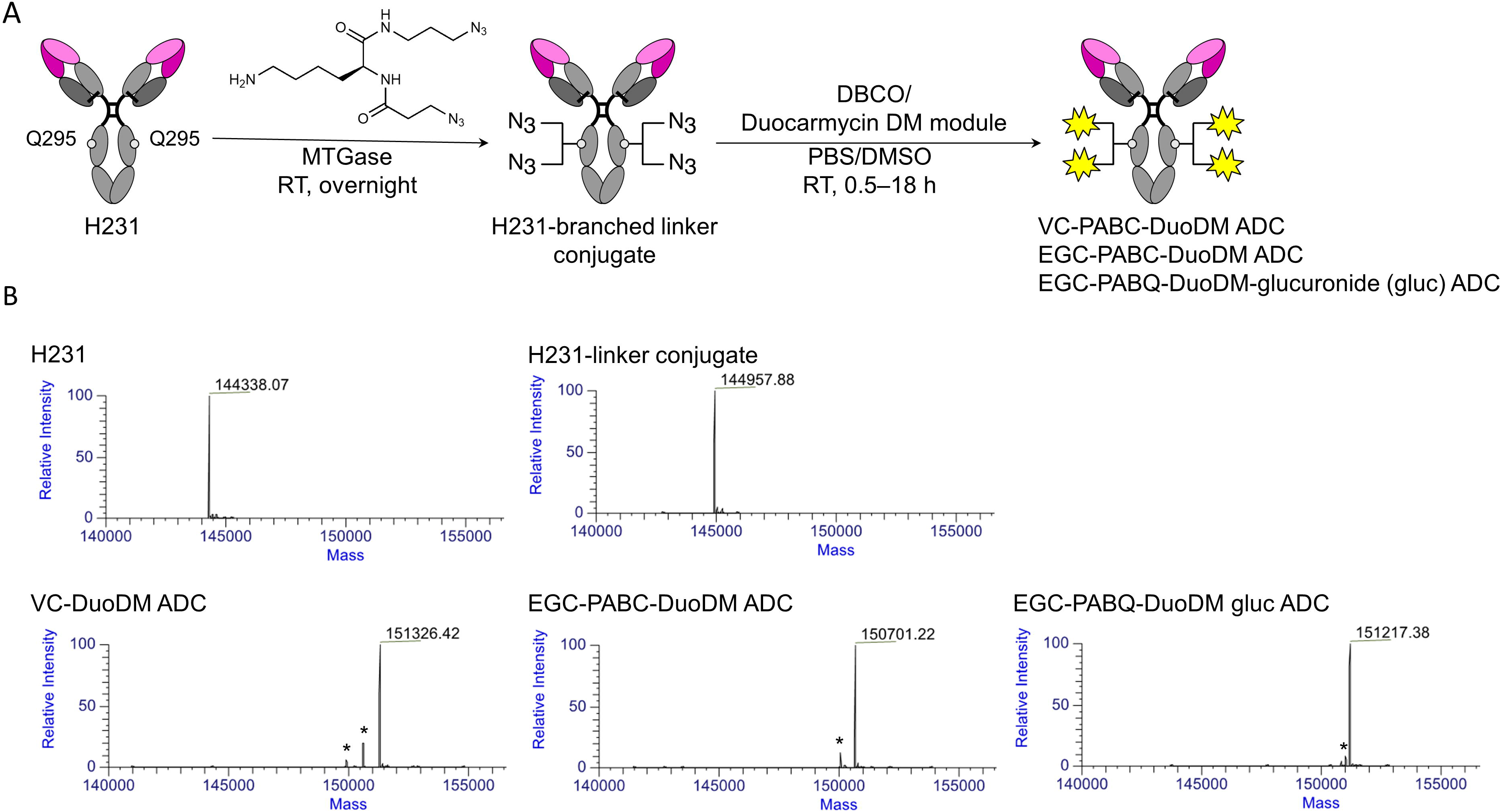
Generation of EREG-targeting antibody-drug conjugates. **A** , Schematic depiction of microbial transglutaminase (MTGase)-mediated conjugation of a branched linker to Q295 of the Fc region of H231 mAb followed by strain promoted alkyne-azide cycloaddition of duocarmycin DM (DuoDM) payload attached to either a cleavable dipeptide valine-citrulline (VC) or tripeptide glutamic acid-glycine-citrulline (EGC) linker to generate H321 ADCs (DAR1=14). **B**, Deconvoluted ESI-mass spectra. First panel, H231 mAb; second panel, H231-branched linker conjugate, third panel, H231-VC-cDuoDM ADC; fourth panel, H231-EGC-cDuoDM ADC; fifth panel, H231-EGC-qDMDM gluc ADC. All homogeneous ADCs have a DAR of 4. *Fragment ions detected in ESI-MS analysis.

### EREG ADCs demonstrate high selectivity and potency in a panel of CRC cells

Prior to evaluating therapeutic efficacy, we measured binding of all three EREG ADCs in parallel in DLD-1 cells. All ADCs demonstrated comparable binding affinities (Kd≈ 2.0 nmol/L; Fig. 5A). We showed EREG ADCs had subnanomolar potency in EREG-high DLD-1 and LoVo cells, whereas unconjugated H231, cmAb, and cADC had no activity (Figs. 5B-C and Supplementary Table S2). CRC cell-killing, rather than senescence, was demonstrated by increased PARP cleavage in DLD-1 and LoVo cells after 48 h treatment with H231 EGC-qDuoDM-gluc ADC (Supplementary Figs. S7E-F). Minimal PARP cleavage was observed with cADC (Supplementary Fig. S7F). Next, we evaluated activity of individual EREG ADCs in CRC cell lines expressing different levels of endogenous EREG and different *KRAS* mutational statuses. Overall, ADC potency based on IC50 and area under the curve (AUC) values for EREG-expressing CRC cell lines was: H231 EGC-qDuoDM-gluc > H231 EGC-cDuoDM > H231 VC-cDuoDM (Fig. 5D-E, Supplementary Fig. S8A-D, and Supplementary Table S2). For EREG-high *KRAS* mutant DLD-1, LoVo, and HCT116 cells IC50 values for EREG ADCs were: H231 EGC-qDuoDM-gluc (IC50s1=10.01-0.0141nmol/L), H231 EGC-cDuoDM (IC50s1=10.06-0.381nmol/L), and H231 VC-cDuoDM (IC50s1=10.12-0.521nmol/L). For EREG-high *KRAS* wildtype SW48 cells, IC50 values ranged from 0.31-0.50 nmol/L. Furthermore, EREG ADCs had minimal effects in EREG-null SW620, RKO, or DLD-1 EREG KO cells, demonstrating efficacy is target-dependent (Fig. 5D-E and Supplementary Fig. S8A-D). These results show EREG ADCs mediate potent cell-killing efficacy in EREG-expressing CRC cells independent of *KRAS* mutational status.

**Figure 5.**
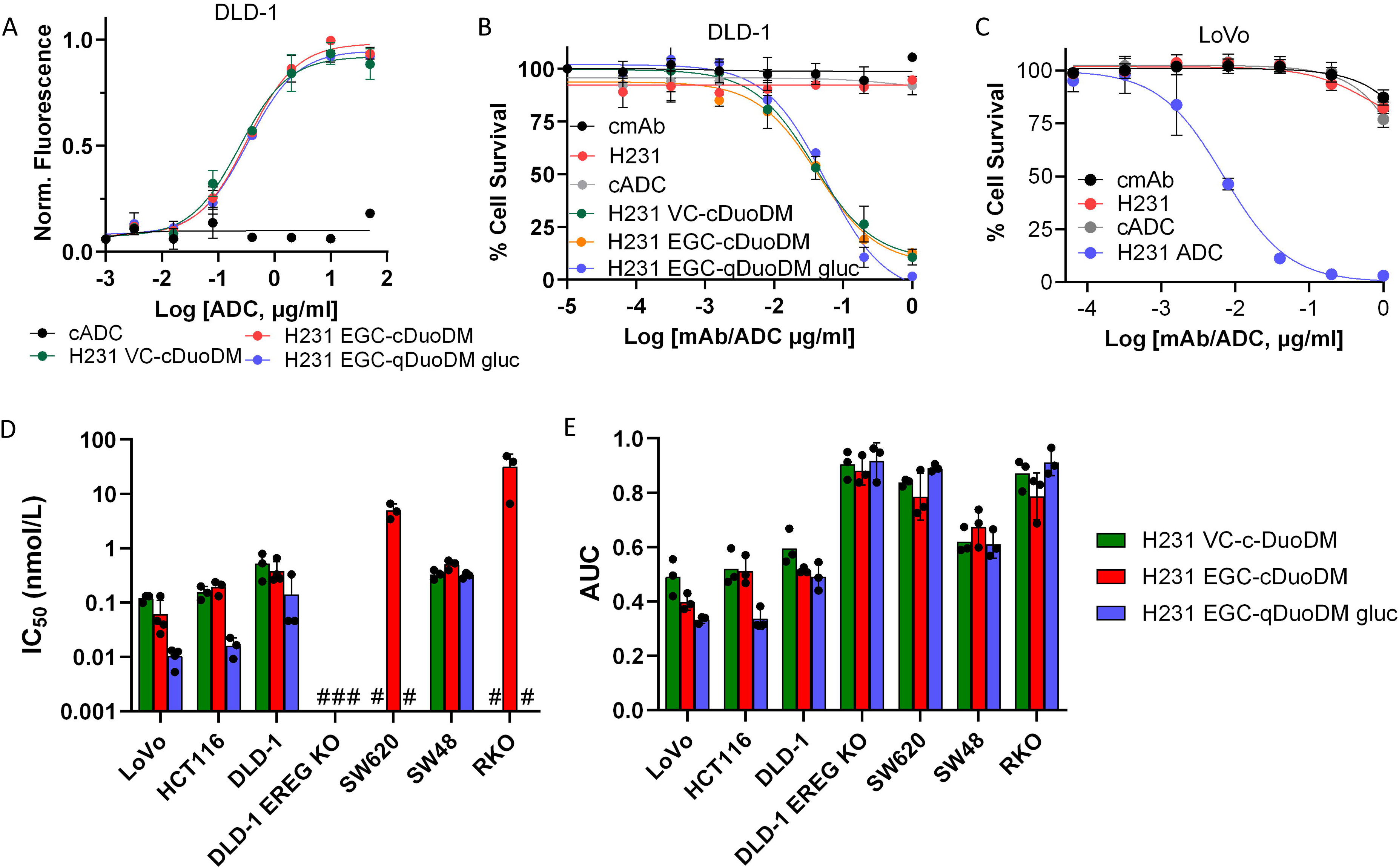
Therapeutic potency and selectivity of EREG ADCs in CRC cell lines. **A**, Binding of H231 ADCs and non-targeting control ADC (cADC, DAR=4) with same linker-payload. **B,** Efficacy of H231 ADCs modified with different linkers, H231, cmAb, and cADC in DLD-1 cells. **C**, Dose-dependent cytotoxicity of H231 EGC-qDuoDM gluc ADC in LoVo cells. H231, cmAb, and cADC have minimal effects. Efficacy of H231 VC-cDuoDM, H231 EGC-cDuoDM, and H231 EGC-qDuoDM gluc in a panel of CRC cell lines with different levels of EREG expression presented as **D**, IC50 values and **E** , Area under the curve (AUC). #, indicates IC50 values cannot be calculated at tested concentrations due to minimal effects. Quantitative data presented as mean ± SD.

### EREG ADCs are well-tolerated and show significant antitumor efficacy in preclinical models of CRC

As H231 binds both hEREG and mEREG, we performed a single-dose escalation and multiple-dose studies to test for ADC on/off-target effects in immunocompetent C57BL/6 mice. Mice were administered vehicle (PBS) or 5 mg/kg or 10 mg/kg EREG ADCs incorporating either VC or EGC linkers and the cDuoDM payload and monitored for weight loss or other signs of overt toxicity. After 2 weeks, mice showed no significant changes in weight, white blood cell (WBC) counts, liver enzymes (alanine and aspartate aminotransferases; ALT or AST), or kidney creatinine (CRE) (Supplementary Fig. S9A-E). As single-dose EREG ADCs were shown to be safe and most FDA-approved ADCs are dosed ≤ 5 mg/kg (37), we performed an additional safety study to examine effects after three 5mg/kg doses of H321 mAb, cADC, or the most potent EREG ADC, H231 EGC-qDuoDM-gluc, administered weekly. Two weeks after treatment was terminated, mice showed no significant changes in weight, WBCs, ALT, AST, or CRE (Fig. 6A-E). Liver and kidney tissues we also collected for H&E staining. As shown in Supplementary Fig. S9F-G, the H231, cADC, and H231 EGC-qDuoDM gluc treatment groups did not have any obvious pathological changes compared to vehicle. These findings suggest EREG ADCs are well-tolerated with no observable on-or off-target toxicities even after multiple doses.

**Figure 6.**
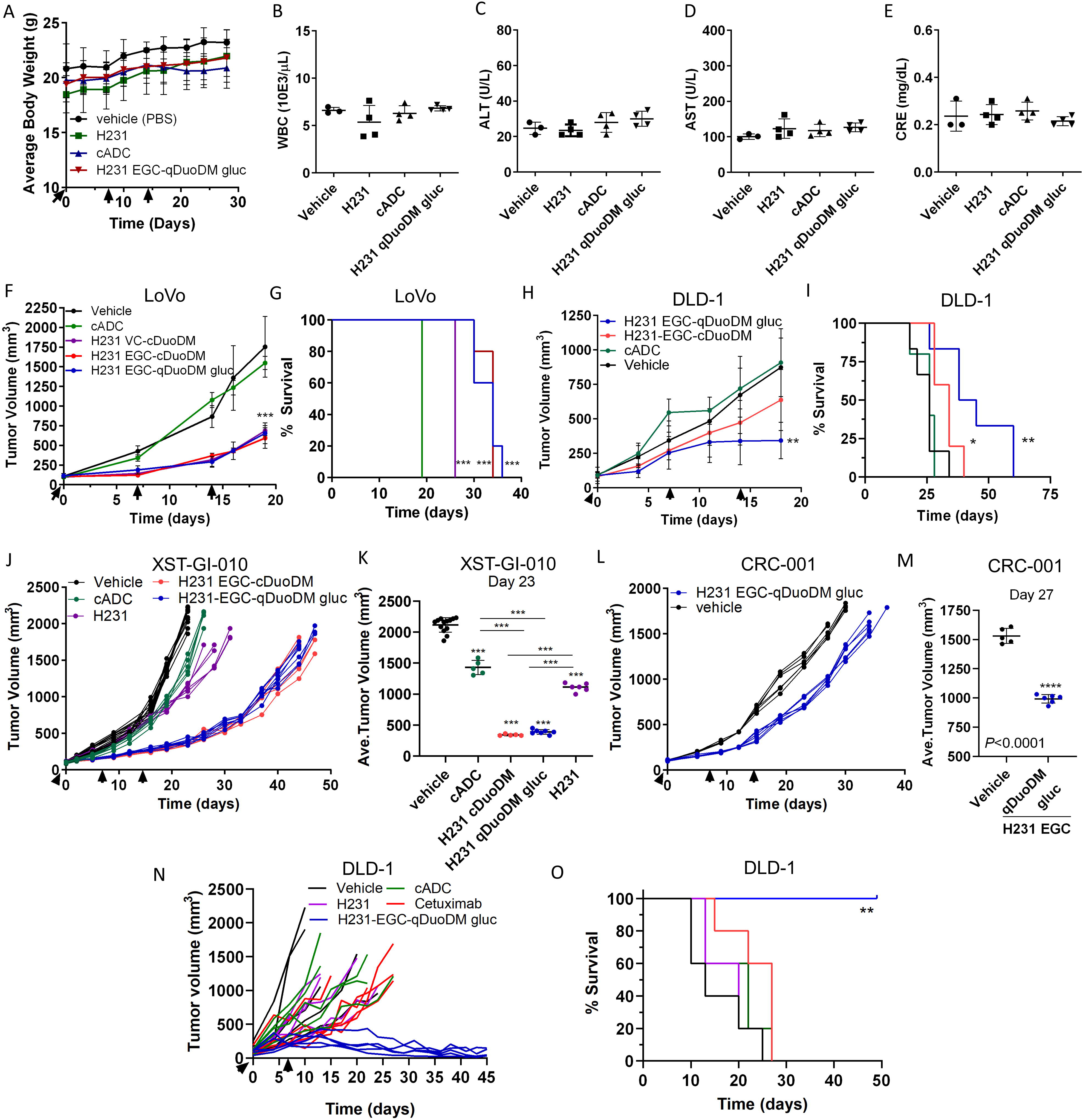
EREG ADC safety assessment and antitumor efficacy in CRC cell line and patient-derived xenograft models. Immunocompetent C57BL/6 mice were treated with 3 weekly 5 mg/kg doses of H231 mAb, cADC, or H231 EGC-DuoDM gluc ADC as indicated, or PBS vehicle. Blood was drawn and mice were euthanized 2 weeks after 3rd dose. **A**, Bodyweight measurements of C57BL/6 mice (n=3, vehicle; n=4 for all other groups). **B**, White blood cell (WBC) counts. **C**, Alanine aminotransferase (ALT) and **D**, aspartate aminotransferase (AST) liver enzyme analysis and E, kidney creatinine in serum. **F**, Antitumor efficacy of EREG ADCs (5 mg/kg) in LoVo nu/nu xenografts (n=9, vehicle; n=5 for all other groups). **G**, Kaplan-Meier survival plot and log-rank test for LoVo xenograft study in F. Vehicle and cADC showed similar survival. **H**, Antitumor efficacy of EREG ADCs (5mg/kg) as indicated in DLD-1 nu/nu xenografts (n=6, vehicle and H231 EGC-qDuoDM; n=5, cADC and H231 EGC-qDuoDM gluc). **I**, Kaplan–Meier survival plot and log-rank test for DLD-1 xenograft study in H. (**J-M**) Antitumor efficacy in patient-derived tumor xenograft (PDX) models. All treatments dosed at 5mg/kg. **J**, XST-GI-010 individual animals (n=13, vehicle; n=5, cADC and H231 EGC-cDuoDM; n=7, H231 EGC-qDuoDM) and **K**, average tumor volume at day 23. **L** , CRC-001 individual animals (n=5, vehicle; n=6, H231 EGC-qDuoDM) and **M**, average tumor volume at Day 27. **N**, Antitumor efficacy of higher dose H231 EGC-qDuoDM gluc compared to cetuximab and control treatment arms in DLD-1 nu/nu xenografts as indicated (n=5/group). All groups were treated with two weekly doses of 10 mg/kg. **O**, Kaplan–Meier survival plot and log-rank test for DLD-1 xenograft study in N. Statistical analysis performed using Student’s unpaired two-tailed t test for two groups or one-way analysis of variance (ANOVA) and Dunnett’s multiple comparison test. Quantitative data presented as mean ± SD. *P< 0.05, **P< 0.01, ***P< 0.001.

Antitumor efficacy was then evaluated in EREG-high *KRAS* mutant MSI-H xenograft models of LoVo and DLD-1 CRC cells (Fig. 6F-I). Mice were administered 5 mg/kg of EREG ADCs weekly for 3 doses, as it was determined safe (Fig. 6A-E and Supplementary Fig. S9). For LoVo xenografts, H231 VC-cDuoDM, H231 EGC-cDuoDM, and H231 EGC-qDuoDM gluc resulted in 64%, 70%, and 70% TGI, respectively, at day 19 when the vehicle cohort reached maximum tumor burden (Fig. 6F). As the tripeptide EGC linkers showed greatest efficacy and increased survival (Fig. 6G), we proceeded with evaluating these 2 ADCs in additional models. In the DLD-1 model, treatment with H231 EGC-cDuoDM and H231 EGC-qDuoDM gluc resulted in 29% and 68% TGI, respectively, at day 18 (Fig. 6H). Importantly, H231 EGC-qDuoDM gluc showed greatest increase in survival in DLD-1 xenograft models (Fig. 6I). We next evaluated EREG ADCs in two EREG-high MSS PDX models of CRC (Fig. 1H): *KRAS* mutant XST-GI-010 and *NRAS* mutant CRC-001. In the XST-GI-010 model, H231 EGC-cDuoDM and H231 EGC-qDuoDM gluc treatment resulted in 88% and 86% TGI at day 23 with similar impact on survival (Figs. 6J-K and Supplementary Fig. S10A). Though unconjugated H231 showed no in vitro activity, treatment in vivo resulted in 50% TGI. As H231 EGC-qDuoDM gluc showed highest efficacy in all models, it was tested in the CRC-001 model and showed 38% TGI and moderate increase in survival (Fig. 6L-M and Supplementary Fig. S10B). Treatment with cADC had no significant impact on tumor growth or survival in CRC cell xenografts, yet had 32% TGI in the XST-GI-010 model (Fig. 6F-J). Analysis of DLD-1, XST-GI-010, and CRC-001 tumors remaining after H231-EGC-qDuoDM-gluc treatment showed reduced EGFR and ERK1/2 phosphorylation and lower expression of EREG compared to vehicle- or cADC-treated tumors, suggesting neutralization of the EGFR pathway (Supplemental Fig. S10C-E). As H231 EGC-qDuoDM gluc showed greatest impact on survival in the DLD-1 xenograft model (Fig. 6H-I), we tested if a higher dosing level would have greater benefit. As shown in Fig. 6N-O, two weekly doses of 10 mg/kg H231 EGC-qDuoDM gluc resulted in tumor regression and significantly increased survival compared to all other treatment groups. H231 and cADC administered at 10mg/kg had minimal effects on tumor growth and survival compared to vehicle. Furthermore, H231 EGC-qDuoDM gluc showed superior TGI and survival relative to the most widely used EGFR-targeted mAb, cetuximab (10 mg/kg). No significant effects in bodyweight or overt toxicity were observed during treatment in nu/nu or NSG mice (Supplementary Figs. S10F-H). These findings demonstrate EREG ADCs have significant efficacy potentially due to combined effects of payload-mediated cytotoxicity and EGFR signaling neutralization in EREG-expressing CRC tumor models irrespective MSI-H/MSS subtype or RAS mutations.

## Discussion

Many clinical reports, including a recent large cohort study of mCRC patients, have validated the prognostic role of EREG expression in *RAS* wildtype patients receiving EGFR-targeted therapies, cetuximab and panitumumab (13–17). While EGFR mAbs only benefit patients with wildtype *RAS*, EREG is highly expressed in both wildtype and mutant *RAS* tumors. Thus, EREG may not only serve as a prognostic marker, but a viable target for the development of novel therapies that benefit patients irrespective of *RAS* or other mutations. Importantly, EREG has considerably low expression in normal human and rodent tissues (4) and *Ereg* KO mice are viable and show no major developmental phenotype (38), suggesting minimal toxicity of EREG-targeted therapies. EREG expression is highest in MSS subtype that typically has worse prognosis and accounts for the majority of CRC. Furthermore, EREG is overexpressed in many other cancers with defined roles in tumor progression including lung, bladder, pancreatic, and breast (5–9). Thus, EREG-targeted therapies could hold benefit to several different tumor types.

In this study, we generated highly-specific EREG mAbs for the development of rationally-designed ADCs for improved CRC treatment. H231 was selected as the lead candidate based on high affinity, species cross-reactivity, and its ability to internalize into cancer cells and traffic to lysosomes, which is critical for payload release (20,21). ImmunoPET and biodistribution studies showed notable uptake of ^89^Zr-labeled H231 by EREG-expressing tumors with minimal uptake in normal non-clearance tissues (Fig. 3). As non-targeting Zr-cmAb remains in circulation for an extended period of time, it can actively perfuse the lungs contributing to the mediastinal imaging signal. Similarly, Zr-H231 is also taken up by the lungs as shown by the ex vivo data, but its high liver and tumor uptake strongly mask signal in other tissues with comparatively lower signal. Clearance of ^89^Zr-H231 was primarily through the liver, which is expected for mAbs. Renal clearance was also higher for ^89^Zr-H231 compared to ^89^Zr-cmAb and could be explained by the elimination of smaller degradation products (i.e., protein fragments) generated by hepatic metabolism of the EREG mAb. Despite the tracer uptake seen on PET, EREG ADC safety studies showed no adverse effects or changes in liver or kidney enzymes in response to different dosing concentrations and frequencies, suggesting they are well-tolerated. Tolerability may also be attributed to the relatively low DAR of our ADCs and selection of a dose comparable to clinically used ADCs to further reduce potential dose-limiting toxicities seen in patients. Previously, an EREG mAb was shown to have significant efficacy in a metastatic model of CRC based on antibody-dependent cell cytotoxicity (ADCC), not growth inhibitory activity (39). Similarly, we found H231 mAb to have no effect on CRC cell survival in vitro, yet moderate antitumor activity in NSG mice. Although NSG mice lack natural killer cells to mediate ADCC, other immune cells (e.g., macrophages and neutrophils) could be involved, though further investigation is needed.

For development of EREG ADCs with enhanced efficacy compared to unconjugated EREG mAb, selection of ADC payload and linker were carefully considered. Typical payloads used in ADC development consist of microtubule inhibitors (e.g., MMAE and maytansines), topoisomerase 1 inhibitors (TOP1i, e.g., camptothecins), and DNA-damaging agents (e.g., duocarmycins and pyrrolobenzodiazepines) (40,41). However, we and others have found colorectal tumors to be relatively resistant to microtubule inhibitors, yet responsive to duocarmycins (34,42). Importantly, trastuzumab duocarmazine (SYD985), an ADC incorporating a duocarmycin payload, demonstrated progression-free and overall survival benefits and is anticipated to be clinically approved for the treatment of HER2 metastatic breast cancers (43). Based on our studies, EREG ADCs are typically more potent in EREG-high CRC cells and neutralize EGFR activity with minimal effects in EREG-low/negative cells. However, potential intrinsic resistance or sensitivity to payload should also be considered in addition to EREG status. For example, though LoVo cells have lower EREG expression compared to DLD-1 cells, EREG ADC is more potent in LoVo cells at 5 mg/kg dose as they are more sensitive to duocarmycins (34). Notably, TOP1is have been effective as chemotherapeutic agents for CRC treatment and, most recently, TOP1i-based ADCs have been clinically successful with the approval of sacituzumab govitecan (Trodelvy) for triple-negative breast cancer and trastuzumab deruxtecan (Enhertu) for tumor agnostic HER2-positive disease and HER2-low metastatic breast cancer (44). Though we selected DuoDM as our payload for first generation EREG ADCs, it is important to note that TOP1is should be considered for future EREG ADC development.

Linkers can control efficacy and stability in systemic circulation and are broadly characterized as cleavable or non-cleavable by which payload release is dependent on lysosomal enzyme cleavage or proteolytic degradation, respectively (45,46). Cleavable linkers have the advantage of bystander effect, meaning released payloads can diffuse and kill neighboring cancer cells to better target tumor heterogeneity. We screened different cleavable linkers: one commercial and clinically validated dipeptide (VC) linker and two tripeptide (EGC) linkers. Consistent with our previous report (26), ADCs incorporating the EGC linker showed enhanced potency in CRC cells in vitro and increased antitumor efficacy and survival in vivo. While ADCs with VC linkers have demonstrated increased risk of adverse effects and low tolerability attributed to premature payload release (47,48), EGC linkers have improved stability against multiple degradation events and potentially broader therapeutic windows (26). The increased TGI and survival of H231 EGC qDuoDM-gluc-compared to H231 EGC-cDuoDM-treated DLD-1 xenografts may then be attributed to its enhanced stability and tumor targeting capacity. Whereas cDuoDM contains an exposed tertiary amine that may foster non-specific binding, qDuoDM-gluc has a covalent bond between PAB and DuoDM that is more cleavage-resistant during circulation and minimizes non-specific binding, as observed in cell-killing at high concentrations in EREG-negative SW620 and RKO cells (Fig. 5D-E and Supplementary Fig. S8A-D). Additionally, β-glucuronidation of qDuoDM-gluc increases ADC hydrophilicity and permits an extra layer of prodrug-based selectivity to avoid prelease of active payload (36,49). It is postulated that only after linker is cleaved by cathepsins in the lysosomes followed by β-glucuronidase conversion of seco-DuoDM to its active form that payload can kill tumor cells and elicit bystander effect to target tumor heterogeneity. Furthermore, increased TGI of H231 EGC-qDuoDM gluc compared to H231 EGC-cDuoDM was observed in DLD-1, but not LoVo tumors. This observation may be attributed to increased endogenous levels of active β-glucuronidase in DLD-1 tumors that convert qDuoDM-gluc prodrug to the active form and/or lower tumor pH, which can further increase β-glucuronidase activity (36). Higher levels of active β-glucuronidase combined with higher specific binding of H231 EGC-qDuoDM gluc compared to H231 EGC-cDuoDM may explain why H231 EGC-qDuoDM gluc is more effective in DLD-1 tumors. Future work will involve elucidating the precise mechanism for increased potency of H321 EGC-qDuoDM gluc. Importantly, we showed two weekly doses of H321 EGC-qDuoDM gluc administered at a higher concentration of 10 mg/kg was well-tolerated, far more potent than cetuximab, and resulted in tumor regression with a significant survival benefit. Notably, we attempted to generate H231 EGC-qDuoDM gluc using mouse in place of human antibody framework for testing the therapeutic window in syngeneic models. However, a technical challenge was encountered due to the presence of additional Q sites in the mouse IgG1 Fc region that were amenable to MTG-based chemoenzymatic conjugation. Thus, site-specific conjugation of mouse IgG1 Fc will require further optimization before EREG ADCs can be evaluated in mouse models with an intact immune system. Furthermore, different dosing concentrations and frequencies should be tested in additional tumor models to determine if EREG ADCs can ultimately eliminate tumors and prevent relapse.

EREG expression positively correlates with expression of another EGFR ligand, amphiregulin (AREG). Similar to EREG, AREG levels predict superior response of mCRC patients to EGFR-targeted therapy (13,15,17). Notably, combined EREG/AREG expression was shown to be associated with progression-free and overall survival among *RAS* wildtype patients treated with anti-EGFR therapy (13,15,17). Another group recently reported the development of an AREG-targeted ADC that showed efficacy in a breast cancer xenograft model (50). AREG ADCs were generated using a mAb that selectively recognizes a neo-epitope of the transmembrane stalk of cleaved AREG. Unlike our EREG ADCs that bind pro-EREG at the tumor cell surface, the AREG ADC did not recognize pro-AREG. Thus, it would be of interest to evaluate how post-translational processing effects these EGFR ligand-targeted ADCs and if co-targeting EREG and AREG provides an improved therapeutic strategy.

Our data show EREG is a promising target for the development of ADCs with broad utility for treating colorectal and other cancer patients with EREG-high tumors. Importantly, EREG ADCs demonstrated therapeutic efficacy with combined payload effects and neutralization of the EGFR pathway in both *KRAS* mutant and wildtype CRC cell lines and tumors, suggesting their potential to benefit a broader patient population than current EGFR-targeted therapies (i.e., cetuximab). As EREG is highly expressed on tumors with minimal surface expression on normal tissues, it will likely have a promising safety profile. As HER2-targeting ADC Enhertu was approved for treating HER2-low breast cancer patients (44), this suggests EREG ADCs may have potential for treating EREG-low CRC patients as well. However, future evaluation in non-human primates and clinical trials are warranted. Recent and exciting progress in validation of EREG as a well-defined biomarker for predicting patient response to EGFR mAbs could be employed to discriminate patients that would respond to EREG ADC therapies, irrespective of tumor MSI or mutational status.

## Authors’ Contributions

Conceptualization and Design: K.S.C; Methodology: K.S.C., J.J., Y.A., K.T., A.A.; Investigation: K.S.C., J.J., Y.A., P.H., Z.L., S.S., S.C.G., S.A., C.G., H.T. J.H.R., A.A., K.T.; Data curation: K.S.C., J.J., Y.A., P.H., Z.L., S.S., S.C.G., S.A., C.G., H.T.; Formal Analysis: K.S.C., J. J., Y.A., P.H.; Writing original draft: K.S.C. and J.J.; Review and Editing: K.S.C, J.J., P.H., C.G., H.T.; Resources: K.S.C., J.H.R., A.A., K.T.; Funding acquisition: K.S.C.; Supervision: K.S.C.

## Acknowledgements

This work was supported by funding from NCI (R01 CA226894 and R21 CA270716) and the Cancer Prevention and Research Institute of Texas (CPRIT, RP210092) to K.S.C., predoctoral fellowships of the Gulf Coast Consortia, on the Training Interdisciplinary Pharmacology Scientists Program (T32 GM139801) to J.J. and P.H., and a predoctoral fellowship in the Biomedical Informatics, Genomics and Translational Cancer Research Training Program (BIG-TCR) funded by CPRIT (RP210045) to S.S. We would like to thank Drs. Vihang Narkar, Pierre McCrea, Qingyun Liu, and Daniel Frigo for helpful project discussions, Drs. Ville Meretoja and Zhengmei Mao for technical assistance, Betty Arceneaux and Dr. Karan Saluja with assistance in patient sample collection and processing, and Martha Thompson for assistance with regulatory approvals for research involving human subjects.

## Conflict of Interest

Y.A. and K.T. are named inventors on all or some of the patent applications (WO2018218004A1, US11629122B2, EP3630189A4 and WO2023122587A3) relating to the linker technologies described in this article. K.T. is a co-founder of and holds equity in CrossBridge Bio. All other authors declare no potential conflicts of interest.

## Notes

### Summary of Updates

Title, authors, and abstract have been updated. Figs, 1, 2, 3, 5, and 6 have been revised and additional data added. Supplementary Figs. S2, S6, S8-10 have been modified. Supplementary Tables S1 and S2 have been modified. Text has been modified to include new data and more details.

