## Supplementary Figures S1-S10 and Supplementary Tables 1-2 for "Antibody-Drug Conjugates Targeting the EGFR Ligand Epiregulin Elicit Robust Anti-Tumor Activity in Colorectal Cancer"

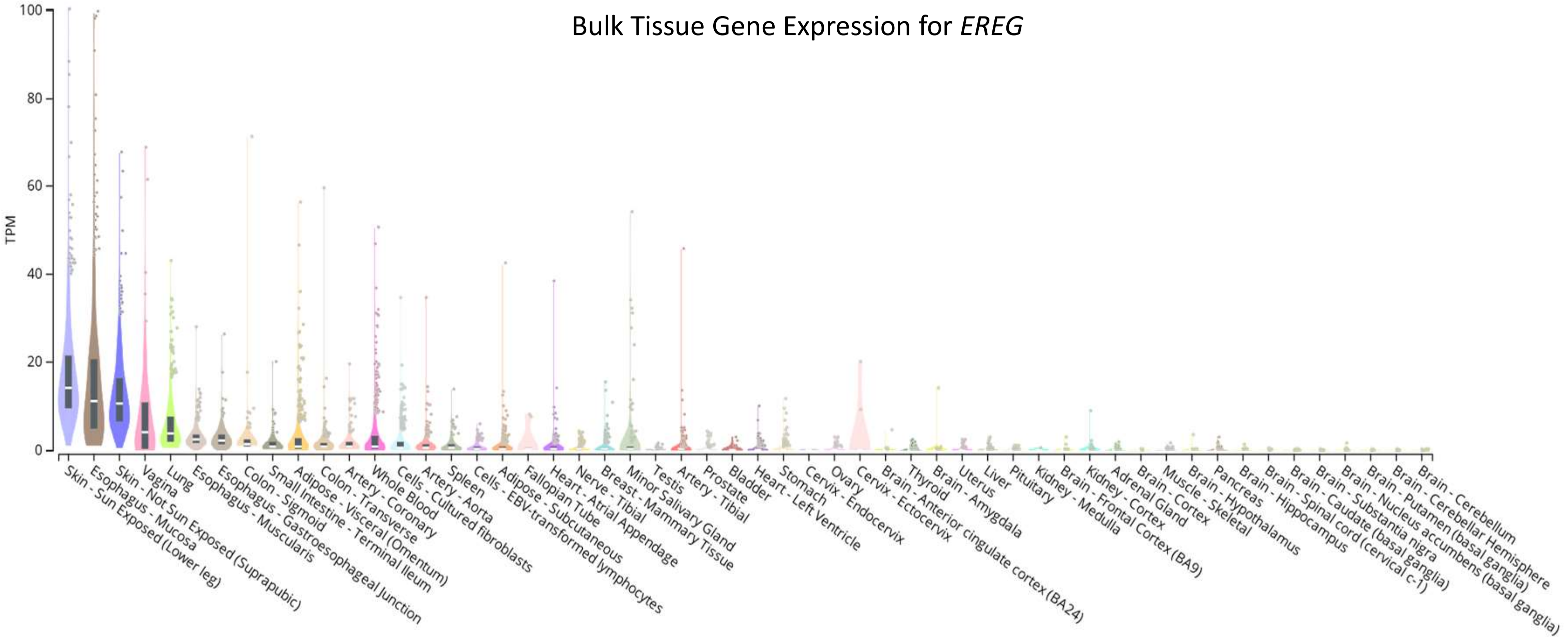


**Figure S1. *EREG* expression in normal tissues and cancer cell lines.** RNA-seq dataset from the Genotype-Tissue Expression (GTEx) project reported as protein-coding transcripts per million (TPM).

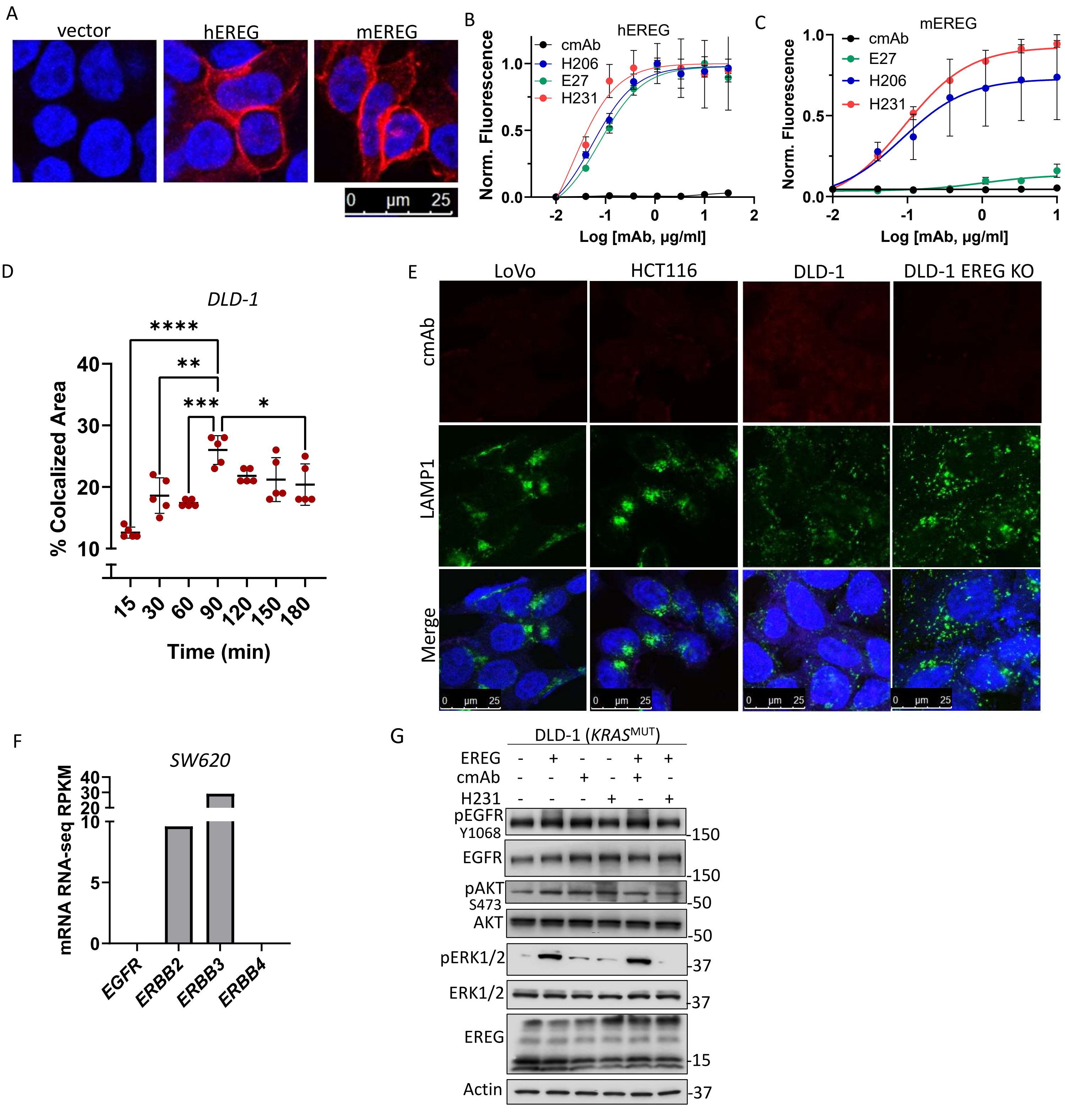


**Figure S2**. **Characterization of EREG mAbs and non-targeting control mAb binding. A**, Immunocytochemistry of vector, hEREG, and mEREG 293T cells using anti-myc-Cy3. **B**, Cell-based binding assays show EREG mAbs H321, H206, and E27 binds recombinant hEREG-293T cells and **C**, H231 and H206, but not E27 bind mEREG-293T cells. Non-targeting control mAb (cmAb) shows no binding in either cell line. **D**, Quantification of H231 (EREG) colocalization with lysosomes at 37 °C in DLD-1 cells over a time-course. Each data point (n=5) represents an image containing 30-40 cells. Statistical analysis performed using ANOVA. Quantitative data presented as mean ± SD. *P< 0.05, **P< 0.01, ***P< 0.001, ****P< 0.001. **E**, Confocal images shows cmAb does not bind or co-localizes with lysosome marker, LAMP1, after 90 mins at 37 °C in EREG-expressing LoVo, HCT116, DLD-1, and DLD-1 EREG KO cells. **F**, CCLE RNA-seq values of HER/ERBB receptors in the SW620 CRC cell line. **G**, Western blot of DLD-1 cells treated in the presence of 15 µg/ml cmAb or H231 for 5 min with or without 300 ng/ml EREG.

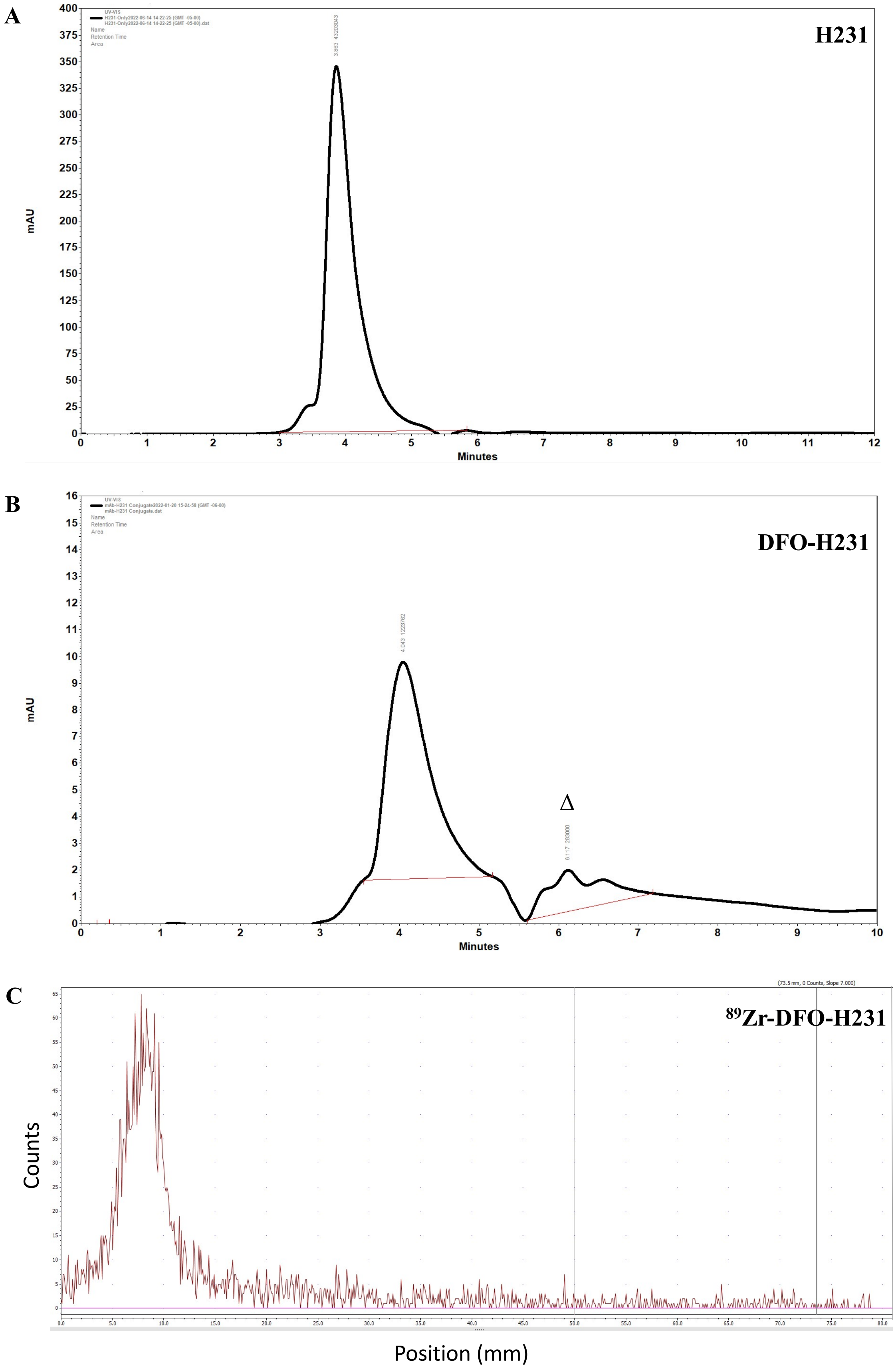


**Figure S3. Characterization of DFO-H231 and $^{89}$Zr-DFO-H231 conjugates.** Representative HPLC analysis showing minimal difference in retention time and peak profiles for **A**, H231 and **B**, DFO-H231. Δ indicates retention time of small molecule impurities under reaction conditions. **C**, Representative radio-ITLC showing high radiochemical purity of $^{89}$Zr-DFO-H231.

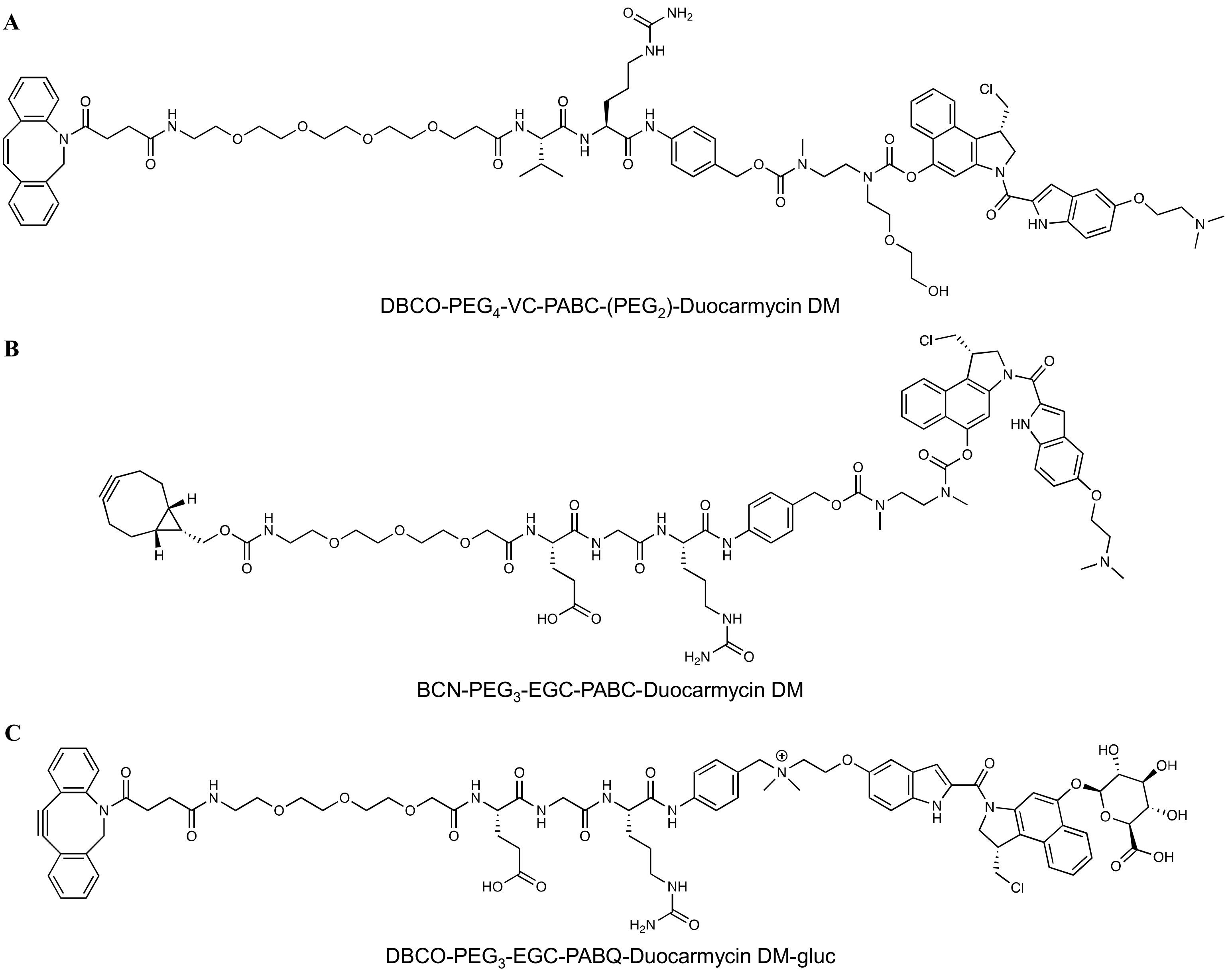


**Figure S4. Chemical structures of duocarmycin-based linker-payloads. A,** DBCO-PEG4-VCit-PABC-(PEG2)-Duocarmycin DM, **B,** BCN-PEG3-EGCit-PABC-Duocarmycin DM and **C**, DBCO-PEG3-EGCit-PABQ-Duocarmycin DM-glucuronide. DBCO, Dibenzocyclooctyne; BCN; bicyclo[6.1.0]nonyne; VC, valine-citrulline; EGC, glutamic acid-glycine-citrulline; PABC, *p*-aminobenzyloxycarbonyl; PABQ, *p*-aminobenzyl quaternary ammonium, and polyethylene glycol (PEG); gluc, glucuronide.

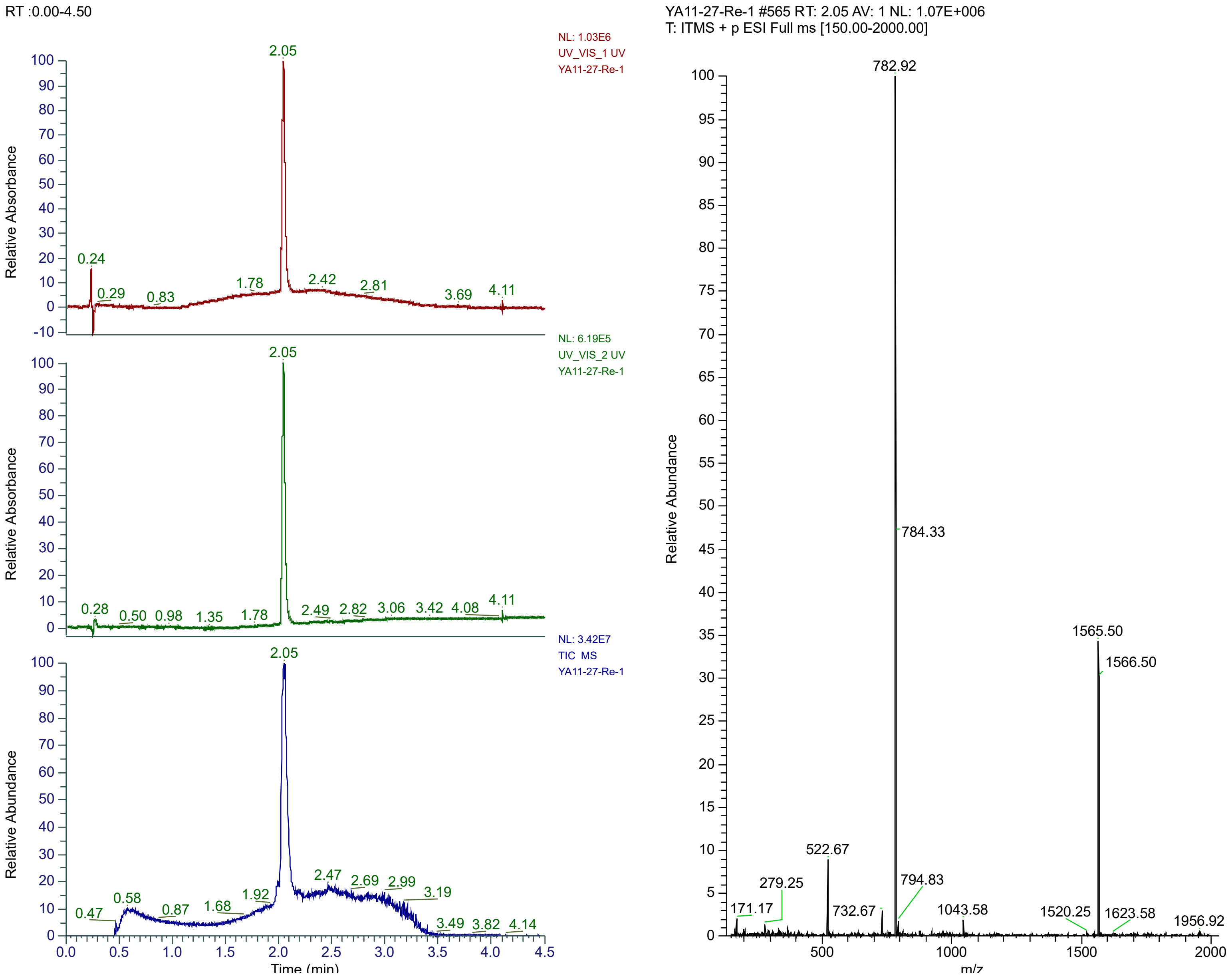


**Figure S5. HPLC and deconvoluted ESI-MS analysis of DBCO-PEG$_3$-EGCit-PABQ-Duocarmycin DM-gluc (purity >95%)**

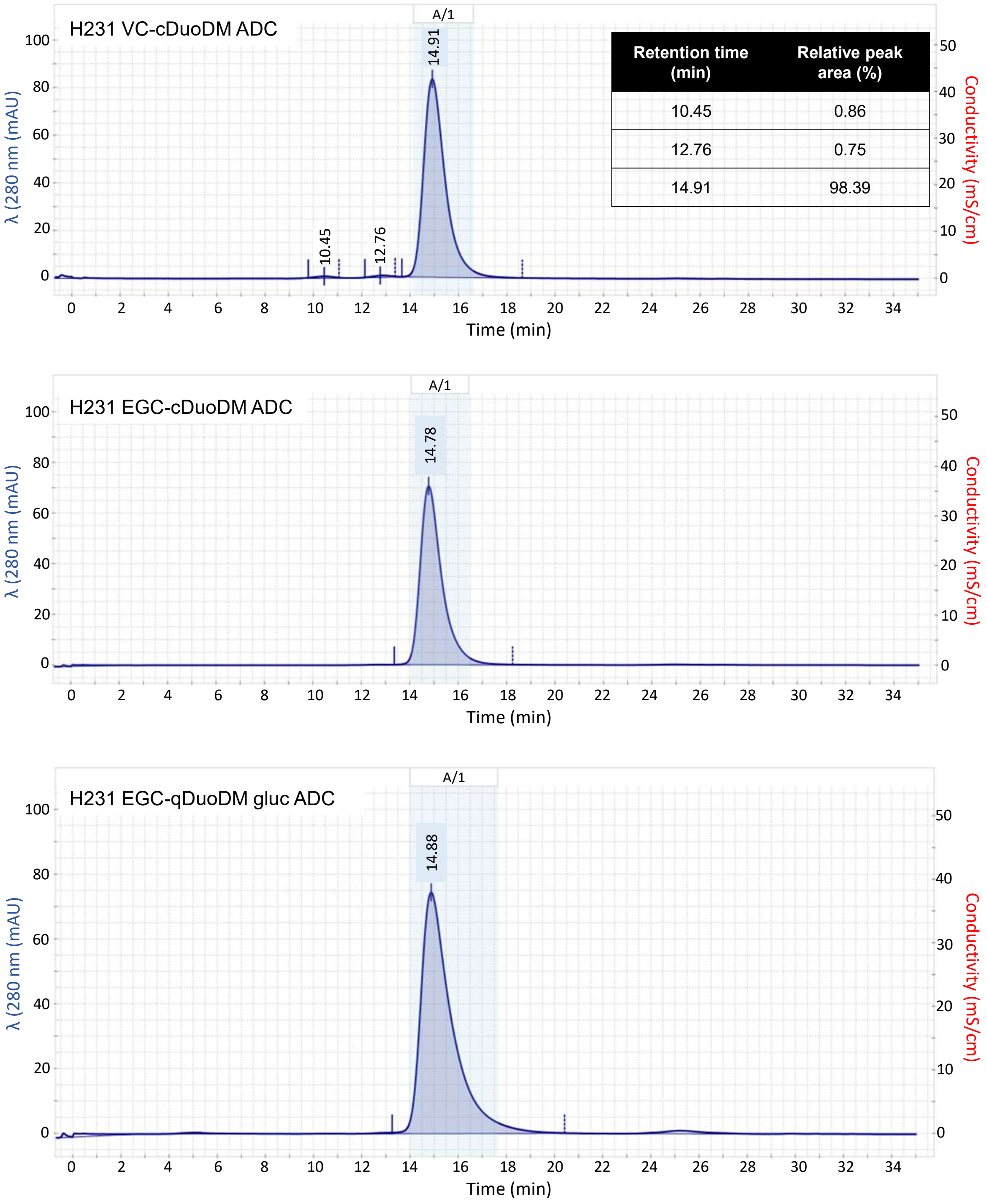


| Retention time (min) | Relative peak area (%) |
|---|---|
| 10.45 | 0.86 |
| 12.76 | 0.75 |
| 14.91 | 98.39 |

**Figure S6. Size-exclusion chromatography analysis after ceramic hydroxyapatite XT (CHT-XT) purification for each of the H231 ADCs.**

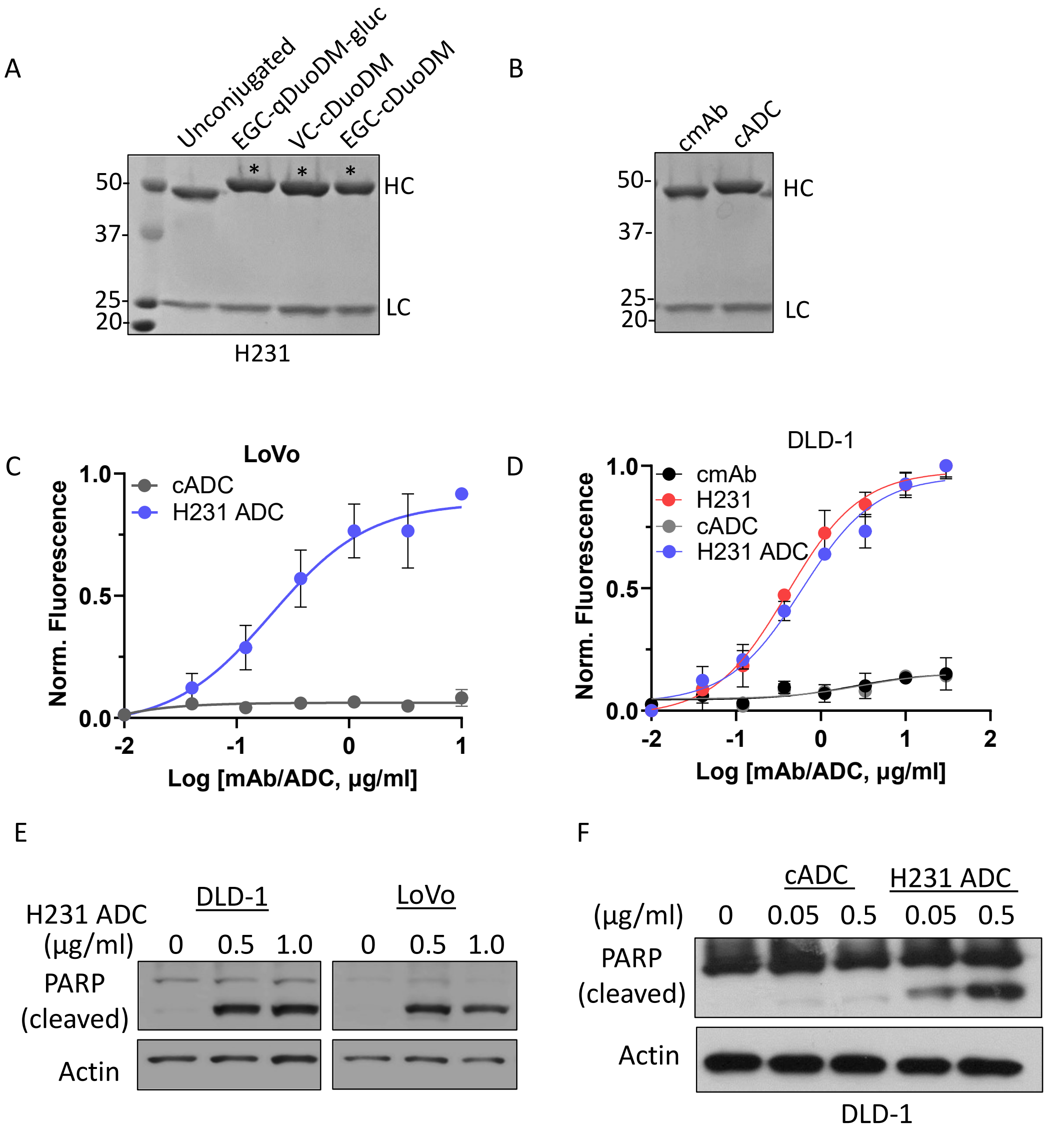


**Figure S7. Characterization of EREG-targeting antibody-drug conjugates. A,** Coomassie blue stained SDS-PAGE of H231 mAb and ADCs and **B,** cmAb and cADC under reducing conditions. *Indicates increased MW of heavy chain (HC) due to conjugated linker-payload on Fc. LC, light chain. **C**, Binding of H231 EGC-qDuoDM gluc ADC compared to control ADC (cADC, DAR=4) with same linker-payload in LoVo cells. **D**, Binding of H231 EGC-qDuoDM gluc ADC compared to unconjugated H231 mAb, non-targeting control mAb (cmAb), and control ADC (cADC, DAR=4) with same linker-payload. **E**, Western blot showing H231 EGC-qDuoDM gluc ADC induces PARP cleavage in DLD-1 and LoVo cells after 2 days and **F**, cADC has minimal effect on PARP cleavage as shown in DLD-1 cells. Quantitative data presented as mean ± SD.

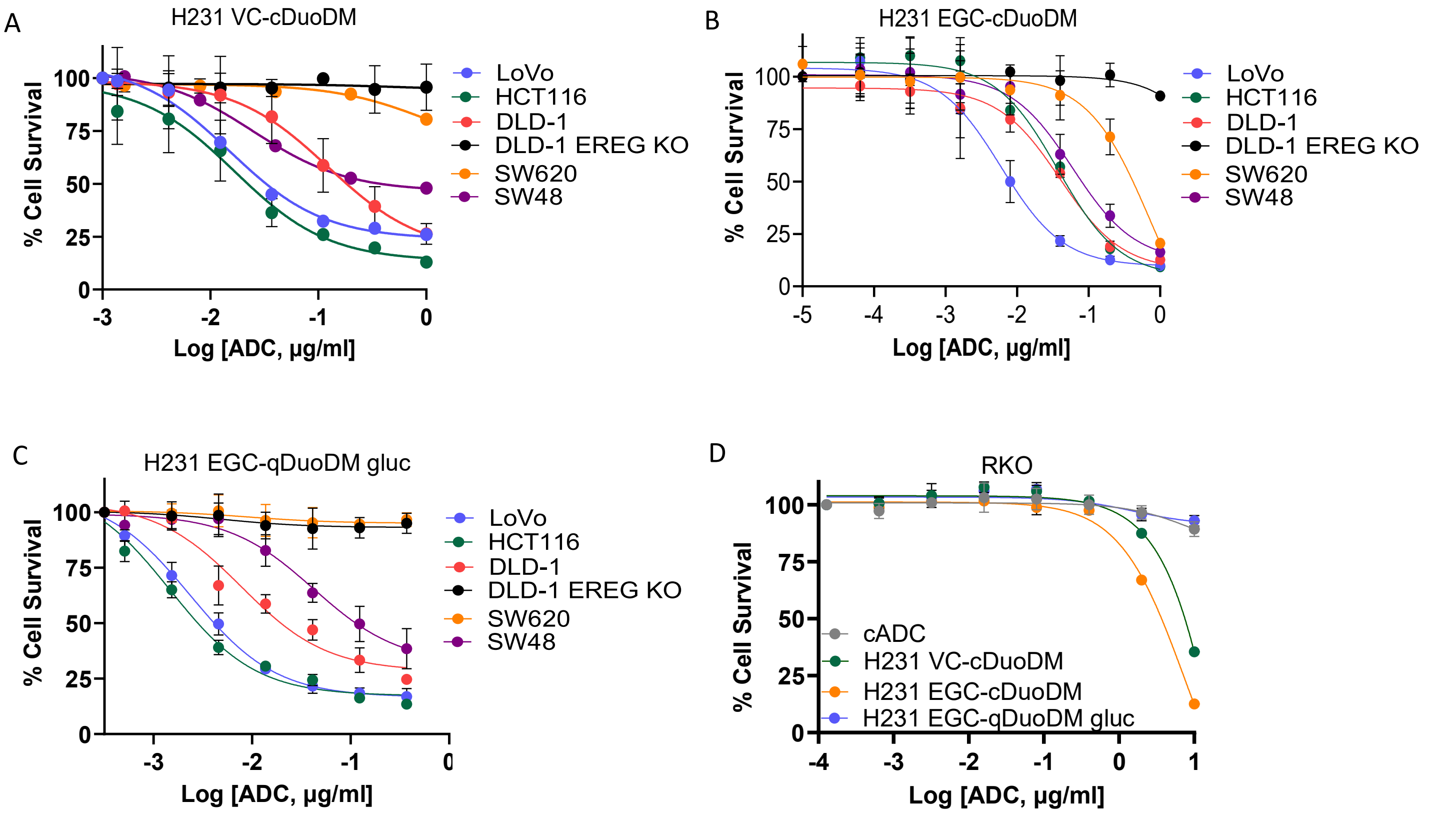


**Figure S8. Cytotoxicity dose-response curves of EREG-targeting antibody-drug conjugates.** Efficacy of **A**, H231 VC-cDuoDM**, B**, H231 EGC-cDuoDM, and **C**, H231 EGC-qDuoDM gluc in a panel of CRC cell lines with different levels of EREG expression. **D**, Efficacy of H231 ADCs modified with different linkers and cADC in EREG-negative RKO cells. Quantitative data presented as mean ± SD.

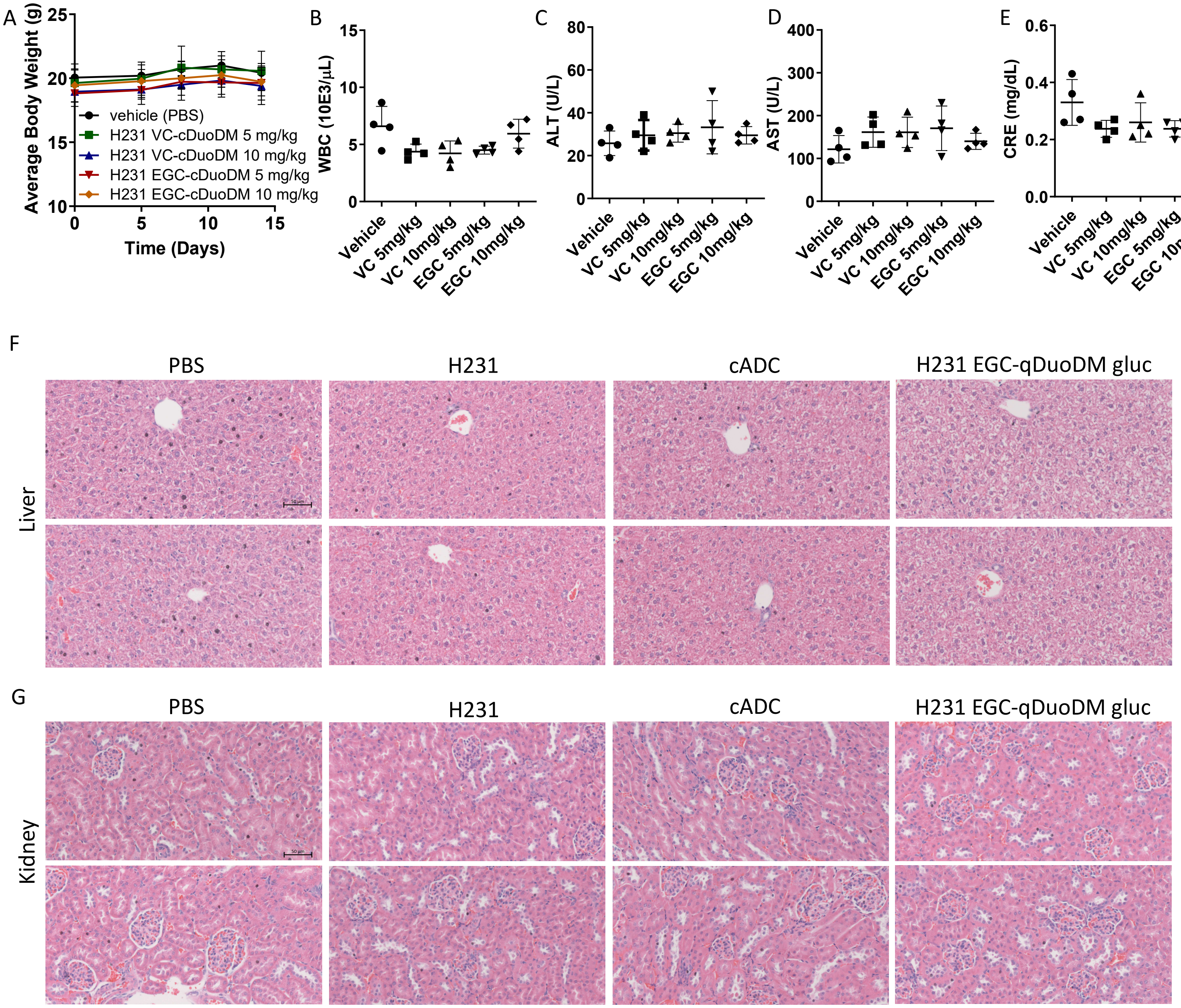


**Figure S9. Safety assessment for H231 VC cDuoDM and EGC cDuoDM ADCs.** Immunocompetent C57BL/6 mice were treated with single-dose EREG ADCs consisting of VC or EGC cleavable linkers as indicated, or PBS vehicle. **A,** Bodyweight measurements of C57BL/6 mice (n=4/group). **B,** White blood cell (WBC) counts on day 14. **C,** Alanine aminotransferase (ALT) and **D**, aspartate aminotransferase (AST) liver enzyme analysis and **E**, kidney creatinine in serum collected from mice on day 14. Statistical analysis was performed using one-way ANOVA and Dunnett's multiple comparison test. Error bars are SD. H&E staining of **F**, liver and **G**, kidney tissues of 2 representative mice from multidose safety study (Fig. 6A- E).

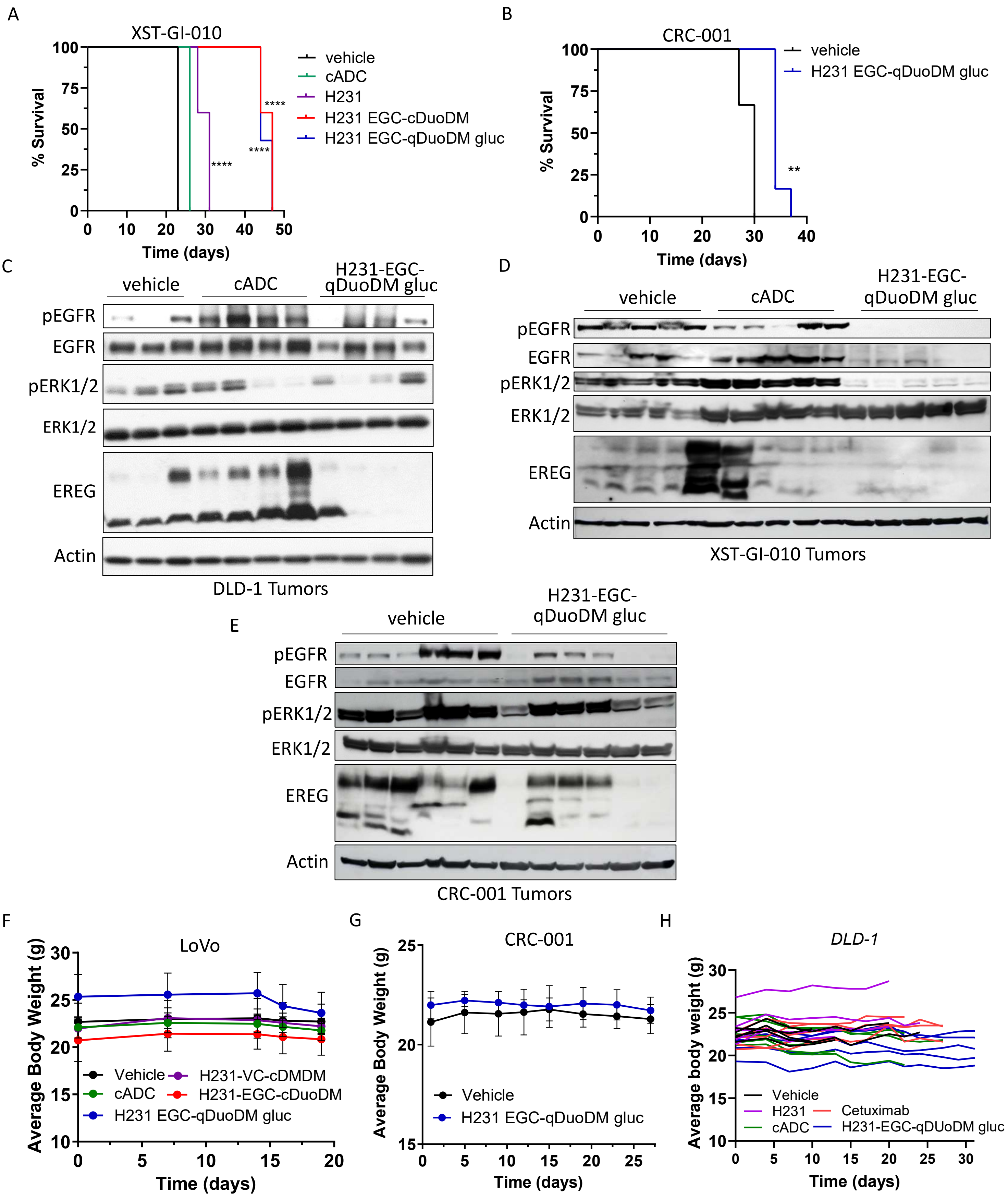


**Figure S10. Biomarker expression and Kaplan-Meier survival plots and bodyweight measurements from efficacy studies.** Kaplan-Meier plot and log-rank test for **A**, XST-GI-010 and **B**, CRC-001 models. $^{**}P< 0.01$, $^{****}P< 0.001$. Western blot analysis of EGFR pathway in tumors from **C**, DLD-1 and **D**, XST-GI-010 after treatment with vehicle, cADC, or H231-EGC-qDuoDM gluc and **E**, CRC-001 after treatment with vehicle or H231-EGC-qDuoDM gluc from Fig, 6H-M after termination endpoints. Average bodyweights for **F**, LoVo nu/nu xenografts and **G**, CRC-001 tumor-bearing NSG mice, and **H**, DLD-1 nu/nu xenografts dosed at 10 mg/kg. Statistical analysis was performed using one-way ANOVA and Dunnett's multiple comparison test. Error bars are SD.

| Cell Line | *RAS*$^{MUT}$ status | PE Fluorescence (Mean) | PE Fluorescence (Range) | Pro-EREG/cell (Mean) | Pro-EREG/cell (Range) |
|---|---|---|---|---|---|
| HCT116 | + | 5079 | 222-15636 | 4988 | 218-15355 |
| DLD-1 | + | 9270 | 483-24412 | 9103 | 474-23937 |
| SW48 | - | 2220 | 163-5471 | 2180 | 160-5373 |
| DLD-1 EREG KO | + | 1186 | 299-3377 | 1165 | 294-3316 |
| SW620 | + | 143 | 16-322 | 141 | 16-316 |

**Supplementary Table S1.** Quantification of surface expressed pro-EREG ligands for a panel of colorectal cancer cell lines.

| Average $IC_{50}$ values (nmol/L ± SD) | | | |
|---|---|---|---|
| **Cell Line** | **H231 VC-cDuoDM** | **H231 EGC-cDuoDM** | **H231 EGC-qDuoDM gluc** |
| HCT116 | 0.15 ± 0.060 | 0.19 ± 0.070 | 0.02 ± 0.009 |
| DLD-1 | 0.52 ± 0.390 | 0.38 ± 0.190 | 0.14 ± 0.160 |
| LoVo | 0.12 ± 0.020 | 0.06 ± 0.048 | 0.01 ± 0.004 |
| SW48 | 0.33 ± 0.093 | 0.50 ± 0.140 | 0.31 ± 0.050 |
| DLD-1 EREG KO | NE | NE | NE |
| SW620 | NE | 4.93 ± 2.79 | NE |
| RKO | NE | 31.66 ± 24.12 | NE |

**Supplementary Table S2.** IC50 values for EREG-targeting H231 ADCs in a panel of colorectal cancer cell lines. NE, no effect. Data represents N=3-4 experiments performed in triplicates.
